# Phenotype-environment matching in ground-nesting birds across and within large-scale biomes

**DOI:** 10.64898/2026.06.08.730407

**Authors:** Javier Medel, Emmanuelle S. Briolat, Silu Lin, Andrew Young, Martin Stevens

## Abstract

Many animals have camouflage appearances that correspond to the habitat where they live, termed phenotype-environment matching. However, this has generally been tested in only a limited number of systems and natural settings, and often not to the relevant vision of key receivers (predators) across different spatial scales. Here, we measured the plumage attributes and putative camouflage used by ground-nesting bird species specialist to particular biomes, specifically tropical rainforest, taiga forest, dry forest, grassland, desert, and tundra from museum specimens. Digital photography and image analysis were used to quantify colour patterns to models of predator vision to understand how different colour patterns may correspond with biome type. With this information, we next created bird models that were photographed *in situ* in the Valdivian temperate rainforest and Patagonian grassland biomes of Chile to quantify the extent to which plumage coloration and pattern traits provide effective camouflage at different spatial scales. In general, we find that specialist ground-nesting birds express a phenotype that better matches the substrate composition and vegetation structure across large spatial scales of their own biome. This study reveals how animal camouflage works across biomes, relative to the visual system of raptor predators, and at the appropriate distance at which detection may occur.

## Introduction

The appearances of animal species show an extraordinary complexity and diversity of colour-pattern combinations. This variation has been associated with different functions, such as camouflage, sexual signalling, and other forms of communication (Darwin, 1859; Wallace, 1889; Cott, 1940), as well as with the mechanisms through which they work (Cuthill, 2019; Merilaita *et al.,* 2017; Stevens & Merilaita, 2011; Fisher, 1915; Zahavi, 1975; Ryan, 1990; Fuller *et al.,* 2005; Endler, 1992). Among these, camouflage is likely the most prevalent use of coloration across animal taxa, and often involves matching key features of the specific patches where the animal lives (e.g. Sargent, 1966; Green *et al.,* 2019), or possessing colour and pattern traits that facilitate concealment in a given habitat, termed phenotype-environment matching (Stevens *et al.,* 2014a; Todd *et al.,* 2006; Nokelainen *et al.,* 2020; A. H. Thayer, 1918). To avoid detection by predators in the environment, a successful phenotype-environment association for camouflage may not only link to the specific background conditions on a small spatial scale (e.g. a moth on tree bark) but also the general characteristics of the local habitat on a larger scale (e.g. forest type and light conditions) (Stevens *et al.,* 2014a; Todd *et al.,* 2012; Nokelainen *et al.,* 2017; Boratyński *et al.,* 2014), particularly for animals with medium to large body sizes (see A. H. Thayer, 1918; Endler, 1978; Stoner et al., 2003a, 2003b; Caro & Koneru, 2021). While many animals are capable of colour change (Duarte *et al.,* 2017; Chiao *et al.,* 2011), others have fixed colours and patterns that need to match the colour, lightness, and pattern of either one background type (i.e. a specialist strategy) or several background types (i.e. compromise or generalist strategy) for effective concealment (Merilaita *et al.,* 1999; 2001; Houston *et al*., 2007; Stevens & Merilaita, 2009; Hughes *et al.,* 2019; 2023). Since most natural environments vary in space and time, this introduces the question of how animal species with fixed colours and patterns match different local backgrounds within the biome they inhabit, as well as the wider scene.

To avoid detection, the appearance of animals often resembles the local substrate composition of their biome. For example, in desert rodents, dorsal appearance closely matches the specific background conditions of their local habitat types (e.g. sand dune or plateau) (Nokelainen *et al.,* 2020), facilitating a phenotype-environment match to the Sahara Desert environment (Boratyński *et al.,* 2014; 2017). Other species such as mice, lizards, toads, vipers and insects also display large phenotypic variation, with dorsal colour tones ranging from light to dark, finely tuned to the specific background characteristics of the substrate types found within their local habitats (Hoekstra *et al.,* 2006; Macedonia, 2001; Marshall *et al.,* 2015; Rosenblum, 2006; Rajabizadeh *et al.,* 2015; Yamamoto & Sota, 2020; Barnett *et al.,* 2021). Similar examples of phenotype-environment matching are found in marine invertebrates that display colour change through plasticity (Stevens *et al.,* 2014a, 2015; Todd *et al.,* 2006; Green *et al.,* 2019; Duarte *et al.,* 2016), as well as in ontogenetic changes in appearance observed in many mammal species (Allen *et al.,* 2011; Caro & Koneru, 2021; Caro, 2005, 2013; Caro *et al.,* 2018; Stoner *et al.,* 2003a; 2003b).

In addition to matching the local substrate composition, an organism’s optimal camouflage is likely to be impacted by the general structure of vegetation of its biome over a wider scale. For example, Endler & Théry (1996) found that in tropical forest-dwelling bird species, the cryptic colour patterns of female and juvenile individuals were similar to random samples of the background in all light environments. Among different biomes, the dark and pale appearance of mammals varies in accordance with the predominant light conditions, which are influenced by features of the vegetation structure. For example, the dark coats of mammal species such as suiforms (pigs and related species) likely match the dark vegetation under low light levels at the ground of dense forests (Caro *et al.,* 2018), while the light coats of desert artiodactyls and lagomorphs are strongly associated with the light coloured arid ground under direct sunlight (Stoner *et al.,* 2003a, 2003b; Caro & Koneru, 2021). Overall, to prevent detection when faced with a variety of different local backgrounds, camouflage patterns should match the biome substrate composition, as well as account for the light conditions influenced by vegetation structure. However, work to date has often focused on phenotype-environment matching at a small scale in line with specific patch types.

Ground-nesting birds are particularly well-suited for large-scale investigations of the relationship between general biome background characteristics and camouflage patterns. Many species rely on camouflage for nest survival (see Loevell *et al.,* 2013; Skrade & Dinsmore, 2013; Stevens *et al.,* 2017; Troscianko *et al.,* 2016; Wilson-Aggarwal *et al.,* 2016; Gómez *et al.,* 2018; Cott, 1940), and inhabit biomes with differing background characteristics. Although ground-nesting birds have been shown to select the most appropriate local backgrounds for egg-laying (see Stevens *et al.,* 2017; Lovell *et al.,* 2013; Alothyqi *et al.,* 2024), many bird species are highly mobile and therefore likely require a certain degree of camouflage against the general background characteristics that feature across their biome.

Most of the evidence suggesting that birds use camouflage to match the general characteristics of their biome background comes from studies primarily focused on trade-offs between signalling and camouflage (Endler & Théry, 1996; Gómez & Théry, 2004, 2007; Heindl & Winkler, 2003). However, they rarely measure the overall bird pattern association and match to the general appearance of the background at large spatial scales, nor evaluate this relationship through the visual perception of their main predators. Consequently, it remains unclear how closely avian camouflage matches the general characteristics of their biome backgrounds versus specific patches.

The background characteristics that birds should match will also depend on predator visual perception and the distance at which their predators are searching for them (see Endler, 1978). For example, certain animals, such as moths, match the fine-scale detail of their feeding or resting substrates (e.g. tree bark) partly because of their small size (Sargent, 1966; Mark *et. al.,* 2022; Walton & Stevens 2018) and because predators such as songbirds are likely to search for them from relatively short distances. Conversely, the medium to large body size of camouflaged birds means that their main predators, such as raptors and mammals (Redpath & Thirgood, 1997; Rollins & Carroll, 2001; Blomberg, *et al.,* 2013; Sargeant *et al.,* 1984) may often scan for them from greater distances (e.g. five to ten metres or more). In this context, the location where birds are most vulnerable to predation becomes more unpredictable compared to animals that occupy specific, fine-scale backgrounds. In short, to avoid detection, ground-nesting birds may need to match their backgrounds at relatively large spatial scales.

Here, we use digital photography and image analysis of both museum specimens and artificial bird models in the field to investigate how the plumage attributes of ground-nesting birds vary with the characteristics of their biome backgrounds, and the extent to which these traits provide effective camouflage when viewed through the visual system of raptors. First, we predict that the plumage colours and patterns of the bird species will differ in a consistent manner between biome types (Table 1). Given that biomes have different environmental characteristics, different bird species inhabiting the same biome type should converge in their phenotype. Second, we expect that ground-nesting birds from different biomes (e.g. grasslands and tropical rainforest) will achieve better camouflage against the substrate composition and the vegetation structure of their own biome than against that of a different biome. Finally, birds should better match their own biome backgrounds over larger rather than smaller spatial scales, in order to avoid the detection of predators that scan for them from greater distances.

**Table 1.** Biome selections regarding general substrate composition, vegetation structure and subjectively judged vegetation complexity.

| Biome | Substrate composition | Vegetation structure | Vegetation complexity |
| --- | --- | --- | --- |
| Tropical rainforest | Wet broadleaf litter, short understory vegetation, and woody debris. Cast shadows from small components (e.g. leaf litter) | Broadleaf evergreen trees with closed canopy and dense understory leads to low light conditions at ground level | High |
| Taiga forest | Wet/dry coniferous needles, woody debris, moss, lichens, and snow. Cast shadows from small and large components (e.g. leaf litter and snow patches) | Coniferous trees with closed canopy and semi/open understory leads to low light conditions at ground level |  |
| Dry forest | Dry/wet deciduous leaf litter, woody debris, and bare ground. Cast shadows from small components (e.g. leaf litter) | Woodland deciduous trees with semi/closed canopy and dense sclerophyll understory leads to moderate to low light conditions at ground level |  |
| Grassland | Dry/wet short grass, shrubs, seasonal flowers, and bare ground. Cast shadows from small and large components (e.g. grass and shrubs) | Tall grass and shrubs standing above the ground leads to high to moderate light conditions at ground level |  |
| Desert | Dry short grass, shrubs, bare ground, pavement, and sand-iron oxides. Cast shadows from small and large components (e.g. stones and shrubs) | Clumps of grass and isolated shrubs standing above the ground leads to high light conditions at ground level | Low |
| Tundra Summer | Wet/dry short grass, shrubs, lichens, rocks, moss, and bare ground. Cast shadows from small and large components (e.g. moss and shrubs) | Shrubs and grass standing above the ground leads to high light conditions at ground level |  |
| Tundra Winter | Snow and cast shadows from snow surface | Snow cover leads to high light conditions at ground level |  |

### Methods Study system

Bird skins were photographed from the collections of the Natural History Museum (Tring, UK), including only ground-nesting bird species considered specialists of tropical rainforest, taiga forest, dry forest, grassland, desert, and tundra that met our selection criteria. First, we restricted our analysis to species specialists of a particular biome, and that incubate their eggs in scrape nests directly on the ground (i.e. without a dome, mound, burrow, or cavity structure). Additionally, we included only species in which incubation is mostly achieved by females, as sexual dichromatism in many bird species is thought to reflect selection for crypsis in females (Gluckman & Cardoso, 2010; Martin & Badyaev, 1996; Gómez & Théry, 2004; Wallace, 1889). Finally, species with migratory movements were excluded, as many migratory species inhabit different biomes (Supporting Information, Table S1). The biomes of interest were selected to cover a wide range of substrate compositions, vegetation structures (i.e. dominant vegetation type), and subjectively judged complexity of vegetation based on the influence of vegetation structure on light conditions at ground level (e.g. high complexity leads to low light conditions) (Table 1).

### Museum data collection

Photography followed a range of past studies (e.g. Stevens *et al*., 2007; 2015; 2017; Troscianko *et al.,* 2016; Troscianko & Stevens 2015; Green *et al.,* 2019; Briolat *et al.,* 2020; Nokelainen *et al*., 2017; 2020), with images taken with a Nikon D7000 converted to a full spectrum / UV sensitivity digital camera with a quartz CoastalOpt UV lens 60 mm (Coastal Optical Systems). For images in the human-visible spectrum, the camera was fitted with a UV-IR blocking filter (Baader UV/IR Cut filter; transmission range: 400-700 nm), and for UV images, a UV pass filter was used (Baader U filter; transmission range: 300-400 nm). To calibrate and linearize the colour channel images (see Image analysis and visual modelling), a ruler and multiple Spectralon standards reflecting light equally at 6.2 % (i.e. dark) and 98 % (i.e. white) between 300 and 750 nm were always placed next to the bird specimen in focus. Following a sequential method as in similar studies (Bergman & Beehner, 2008; Stevens *et al.,* 2007) all photographs were taken using the same camera settings (i.e. f/8, ISO 200, RAW format), lighting conditions, and at a similar distance. To reduce the effects of directional lighting from windows and museum lamps, and to produce a spectrum similar to D65 daylight conditions, an Eye Colour Arc Lamp MT70D was used as the primary light source.

### Image analysis and visual modelling

To linearize and calibrate the images of museum bird skins, the camera response was standardized using the grey standards (see Stevens *et al.,* 2007; Troscianko & Stevens, 2015). All images were aligned using an automated script with the Image Calibration and Analysis toolbox-MicaToolbox (see Troscianko & Stevens, 2015), in ImageJ (Schneider *et al.,* 2012). This process was applied to the longwave (LW, “red”), mediumwave (MW, “green”), and shortwave (SW, “blue”) channels of the visible spectrum, as well as the ultraviolet (UV “ultraviolet”) channel. For the visual model system, the peafowl (*Pavo cristatus*) was used as a model for the violet-sensitive (VS) class of colour vision in birds (Hart, 2002), as seen in raptors (Ödeen & Håstad, 2003).

### Identification and selection of plumage attributes

To objectively identify different plumage regions for analysis, a thresholding method was applied to the images to separate areas of colour and marking attributes on the bird skins. Calibrated images were converted to a binary format using a standard threshold value, where black pixels (value = 0) represented markings and white pixels (value = 1) represented the background plumage colour (see Stevens, 2011; Stoddard & Stevens, 2010). The threshold value was determined based on the appearance of the reflectance standards: when the white area of the standard turned completely white, without any black pixels. Once the background plumage coloration was identified, specific patches from that area were selected. For the dorsal area, these included the back (e.g. mantle), wing (e.g. coverts and secondary feathers of only one wing) and rump. With respect to the ventral area, the breast plumage patch was selected (Supporting Information, Fig. S1).

### Colour analysis

To analyse how the colour and luminance of the dorsal and ventral areas vary among bird species, three colour metrics were calculated for each plumage patch: hue (i.e. the actual colour type, such as red or blue), saturation (i.e. the amount of a certain colour compared to white light), and luminance (i.e. perceived lightness). To calculate hue and saturation, each cone catch value was standardized as a proportion of the total response across the four single cone types (i.e. LW, MW, SW, and UV) (Endler & Mielke, 2005; Kelber *et al.,* 2003; Stoddard & Prum, 2008). Using this approach, saturation was measured as the Euclidean distance of the colour patch from the achromatic centre in colour space, with larger values indicating greater colour richness within a specific region of the spectrum (see Stoddard & Prum, 2008).

Hue was quantified based on a ratio of reflectance in different parts of the spectrum (i.e. LW, MW, SW, and UV) representing the relative stimulation of photoreceptors (Komdeur *et al.,* 2005; Spottiswoode & Stevens, 2011; Stevens *et al.,* 2014a; 2014b). Broadly, this approach follows methods developed in previous studies that aim to quantify colour perception as encoded by putative opponent colour channels in visual systems (see Komdeur *et al.,* 2005; Spottiswoode & Stevens, 2011). Because it is unknown which opponent colour channels birds have, following previous studies a principal component analysis (PCA) was used to determine which colour channels to construct, based on a covariance matrix of the standardized cone photon catches derived from the avian visual model (see Spottiswoode & Stevens, 2011; Komdeur et al., 2005). Because PC1 explained most of the variance, it was used to construct the colour channel being calculated as: hue = (LW) / ((UV+SW+MW) / 3). This equation contrasts long wavelength colours (LW) against ultraviolet (UV), short wavelength (SW), and medium wavelength (MW) colours. Therefore, higher hue values indicate that the plumage patches of specialist ground-nesting bird species are relatively redder (e.g. see Briolat *et al.,* 2020). Finally, luminance was measured using double cone values derived from the avian visual model (Osorio & Vorobyev, 2005; Jones & Osorio, 2004).

### Pattern analysis: Granularity

A granularity analysis was used to calculate the pattern size and contrast of ventral and dorsal plumage patches among specialist ground-nesting bird species, following similar methods used to measure egg pattern mimicry in cuckoos and pattern attributes for camouflage in cuttlefish (Barbosa *et al.,* 2008; Hanlon *et al.,* 2009; Stoddard & Stevens, 2010). This analysis decomposes calibrated images by applying a Fast Fourier transform (FFT) and multiple band-pass filters (Godfrey *et al.,* 1987) enabling the identification of markings of different sizes (Barbosa *et al.,* 2008; Stoddard & Stevens, 2010; Stevens et al., 2014). A log scale arrangement was applied, starting at two pixels for the smallest markings and increasing by a factor of two up to a maximum of 1,024 pixels for the largest markings.

From the granularity spectrum, several metrics of plumage patterning were calculated. First, the dominant marking size, which corresponds to the location of the maximum energy value among the overall spectra. Next, the proportion energy, a measure of pattern diversity, or how much one pattern size dominates: higher values indicate that the pattern is dominated by a particular marking size. Finally, the total energy, which provides a measure of the marking contrast, with higher values indicating greater contrasting markings overall (Chiao et al., 2009; Stoddard & Stevens, 2010).

### Field experiments

To test the effectiveness of plumage attributes for camouflage within and between biomes, we photographed bird models that resemble the appearance of specialist ground-nesting bird species from two focal biomes, grassland and tropical rainforest (Fig. 1). The Valdivian temperate rainforest was selected as a substitute for tropical rainforest, which does not occur in Chile, as its complexity of vegetation structure is similar to a tropical rainforest vegetation (see Alaback, 1991; Darwin, 1839, p. 271). For the temperate rainforest, fieldwork took place in Parque Oncol and Reserva Natural Pilunkura, both private protected areas near Valdivia (centred on 39°48′51″S 73°14′45″W) in the Los Ríos region, between 22 February – 29 March 2020, Chile. For Patagonian grassland, photographs were taken in Torres del Paine National Park, and the private protected area Leona Amarga, near Puerto Natales (centred on 51°43′35″S 72°30′22″W) in the Magallanes and Chilean Antarctic region, between 2 February – 16 March 2020, Chile. Fieldwork was conducted during the austral summer season (Supporting Information, Table S2). The study was originally intended to include the Atacama Desert as a third biome, but this was not possible due to COVID pandemic restrictions. All work was approved by the University of Exeter Animal Ethics Committee (application number eCORN001817) and carried out with the permission of landowners and protected area managers.

**Figure 1.**
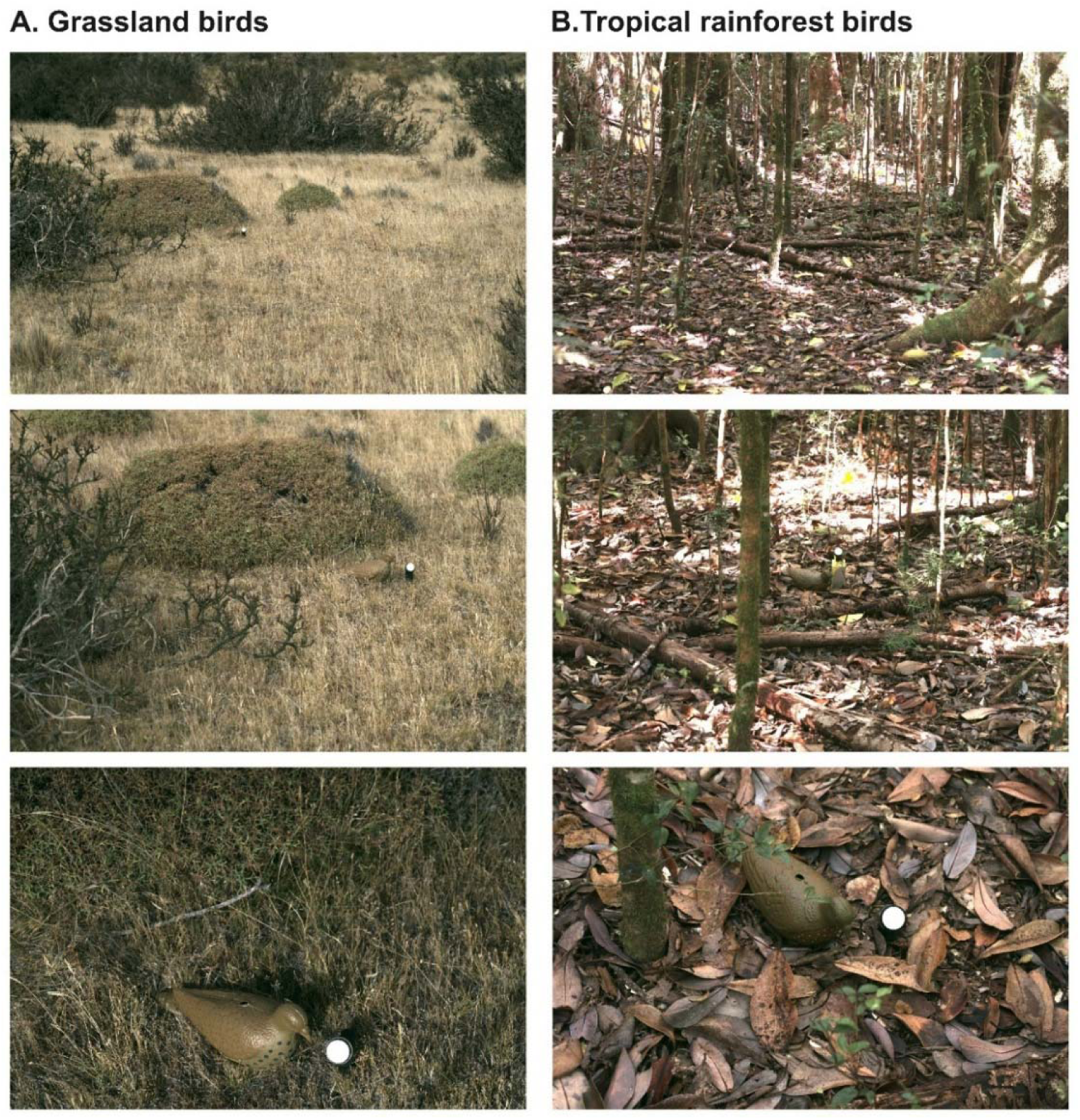
Images of bird models across different spatial scales of biome backgrounds. A) Grassland bird: This shows the mean treatment of grassland bird models across different spatial scales of the Patagonian grassland biome. B) Tropical rainforest bird: This shows the 65% treatment of tropical rainforest bird models across different spatial scales of the Valdivian temperate rainforest biome. As the photograph distance increases, the area in focus near the bird model includes progressively wider characteristics of the substrate composition and vegetation structure of the biome.

### Bird model design

Bird models were created by painting plastic pigeon decoys (A1 Decoy, Chestnut House, Chesley Hill, Wick, Bristol, UK). Plumage colour, marking size, and marking luminance measurements were determined from the general plumage attributes across five individuals from each of 11 tropical rainforest bird species and 22 grassland bird species, resulting in a total of 55 tropical rainforest and 110 grassland bird specimens. To address whether bird models better match the backgrounds of their own biome, we first made an average bird model per biome, each reflecting the mean values of plumage attributes of tropical rainforest and grassland birds (hereafter ‘mean bird models’). These bird models were photographed both in their own biome and the other biome, with the aim to investigate if bird models of different biomes have a better match to their original vegetation structure and substrate composition rather than to a different one. However, it is important not to base our results on only one bird model per biome type, and so we additionally created four further model treatments for each grassland and tropical rainforest biome. Each treatment was painted according to a different quantile value (75%, 65%, 35%, and 25%) of the plumage attributes of interest, derived from measurements of original bird skins (hereafter “quantile bird models”). These quantile bird models were used to introduce variation in the camouflage appearance and to reflect the natural variation in plumage attributes observed among bird skins of specialist ground-nesting bird species within a given biome. This approach ensured that bird models within each treatment were not identical but instead represented variation around the mean bird model appearance. In total, five bird models were created per grassland and tropical rainforest biome (Supporting Information, Fig. S2).

To identify paints that matched the plumage colour and marking luminance metrics of the museum bird skins, 320 colour cards from various paint brand companies were photographed. Quantification of colour and luminance contrast between museum bird skins and paint colour card cone catches was conducted using the log form of the Vorobyev & Osorio (1998) receptor noise model, considering the visual perception of avian predators. To account for receptor noise, a Weber fraction value of 0.05 was applied for the most frequent cone type, based on values from other vertebrates (Vorobyev & Osorio, 1998; Vorobyev et al., 1998). To calculate avian predator perceived chromatic contrast, the relative proportions of cone types in the peafowl (*Pavo cristatus*) were used (LW = 0.92, MW = 1.00, SW = 0.81, VS = 0.54; Hart, 2002). To quantify luminance, we used double cone values derived from the avian visual system (Hart *et al.,* 2000; Jones & Osorio, 2004; Osorio & Vorobyev, 2005) in the achromatic version of the Vorobyev & Osorio (1998) model. Just-noticeable differences (JNDs) calculated from photon catch cone values (Vorobyev & Osorio, 1998) were used to assess the degree of matching between paint colour cards and the plumage attributes of museum bird skins. Colours with JND values below one are generally considered to be indistinguishable, with low JND values likely still difficult to differentiate except under perfect lighting conditions, and higher values indicating increasingly likely colour discrimination (Siddiqi *et al.,* 2004). Overall, the most accurate colour and luminance matches between colour cards and museum bird skins had low JND values, suggesting that the bird models were generally difficult to distinguish under typical conditions.

Granularity analyses (see Chiao et al., 2009; Stoddard & Stevens, 2010) were used to calculate the dominant marking size from the dorsal and ventral plumage patches of museum bird skins. From the granularity analyses, the dominant marking size was detected as the pixel size with the maximum energy (see Pattern analysis: Granularity). To paint the dominant marking size on the bird models, the pixel values were transformed to millimetres. The ‘Batch Scale Bar Calculator’ tool in ImageJ was used to identify a minimum standard pixel value per millimetre. A stencil was used to paint circular markings of the correct size onto the corresponding bird models. The dominant marking size was painted with its respective luminance colour value from the JND analysis.

### Bird model photography in biomes

Field photographs were taken with the same set-up as used for photographing bird museum specimens, except that a Spectralon™ reflectance standard with a larger surface area (Labsphere, Congleton, UK) was used, reflecting light equally at 40 per cent between 300 and 750 nm (Fig. 1). Following image processing, each bird model and its respective background were selected in ImageJ (Supporting Information, Fig. S3).

The same model of visual discrimination (Vorobyev & Osorio, 1998) used to match the colour cards and the museum bird skins was subsequently used to examine how visible the colour and luminance attributes of bird models were for raptor perception against natural backgrounds. Similarly, the same granularity analyses (Chiao *et al.,* 2009; Stoddard & Stevens, 2010) used to calculate the dominant marking size from the museum bird skins were used to calculate the dominant marking of natural backgrounds at different spatial scales. In general, this approach followed the same granularity analysis of the multispectral images.

### Statistics

All statistical analyses were conducted in R v4.1.2 (Schulz, 2024). Linear mixed-effects models were implemented using the lme4 package version v1.1Ö28, applying maximum likelihood estimation (Bates *et al.,* 2015). The Kruskal-Wallis and Welch ANOVA tests were run using the stats package v4.0.5 (R Core Team, 2021). To check for normality, square root transformation, inspection of regression plots, and normality tests were used. Non-parametric tests were used when the data could not be transformed to fit these assumptions.

### Prediction: plumage attributes of specialist bird species will differ in a consistent manner between biomes

Apart from the proportion energy of the dorsal and ventral plumage patches and the marking size of dorsal plumage patch, normality tests and analysis of residuals showed that all colour and pattern metrics were non-normally distributed. A logarithmic and square root transformation was performed, but the data did not fit a normal distribution. To check for homoscedasticity, a Levene’s test was performed; the data showed heteroscedasticity or non-homogeneous variance in most of the cases, except for marking size of the ventral and dorsal plumage patches. For colour metrics, a Kruskal-Wallis test was used to analyse how ventral and dorsal plumage patches of ground-nesting bird species differ among biomes. To improve the visualization of hue data on the graphs, hue was standardized to facilitate easier illustration of results relative to other colour metrics by dividing each value by the maximum hue value of the data set. For pattern metrics, we used a Kruskal-Wallis test to analyse how total energy of dorsal and ventral plumage patches and marking size of ventral plumage patch in ground-nesting bird species differ among biomes. For proportion energy of dorsal and ventral plumage patches, and the marking size of dorsal plumage patch, we used a Welch ANOVA test instead of a one-way ANOVA because it accounts for unequal variances.

### Prediction: bird models will achieve better camouflage against the substrate composition and the vegetation structure of their own biome than against that of a different biome

A normality test and analysis of residuals showed that most of the data were non-normally distributed, except for JND values from ventral colour, marking luminance, and size among tropical rainforest bird models at different spatial scales. The same analysis showed non-normal distributions for dorsal coloration and for ventral marking luminance in mean bird model treatments among different biomes. In these cases, square root transformation of JND values best fitted the assumption of normality of error. Only in the case of marking size attributes of bird models did the data not fit a normal distribution. Levene’s test was performed to check homoscedasticity, and the data showed homogeneous variance or very low levels of heteroscedasticity, except for marking size among mean bird model treatments in the temperate rainforest biome.

Linear mixed models (LMMs) were used to test the level of matching in colour and marking luminance (i.e. square root transformed JND data) of bird models to their own and different local backgrounds across biomes. The model included an interaction between ‘Source Birds’ (whether the bird models were created from the characteristics of tropical rainforest or grassland birds) and ‘Photographed Biome’ (whether bird models were photographed in temperate rainforest or grassland biome). These two terms and their interaction were therefore set as the key fixed effect predictor variables of interest, with background site location and distance fitted as random effects (e.g. Mulder *et al.,* 2021; Stevens *et al.,* 2017). Hereafter, by specifying background site location as a random factor and biome as a fixed effect, we controlled for pseudoreplication in all statistical models. This was necessary because comparisons of camouflage matching to different backgrounds were based on large matrices in which each bird model was compared with its own and different background locations from both biomes: temperate rainforest and grassland (e.g. Stevens *et al.,* 2017). An example of the full mixed model structure was therefore: lmer(JNDdata ∼ SourceBirds * PhotographedBiome + (1|BackgroundSiteLocation) + (1|Distance)). These analyses were restricted to data from the ’mean bird’ models, as only these were photographed across both target biomes.

### Prediction: bird models should better match their own biome backgrounds over larger rather than smaller spatial scales

A linear mixed model was also used to test the level of matching between the colour and marking luminance (i.e. square root transformed JND data) of all bird model treatments (‘mean bird’ and ‘quantile birds’) to their own biome across different spatial scales of the backgrounds. We ran LMMs on each colour metric (i.e. colour and luminance) as the response variables with distance (i.e. 1.25 m, 5 m, and 10 m) set as the predictor variable of interest, with bird model treatments and background site location included as a random effect (e.g. Stevens et al., 2017). The analyses were conducted separately and without an interaction: we performed one set of distance analyses for the grassland birds against their own biome (i.e. grassland), and another set for the rainforest birds against their own biome (i.e. temperate rainforest). Thus, the mixed model was structured as follows: lmer(JNDdata∼Distance + (1|BirdModelTreatment) + (1|BackgroundSiteLocation)). For pattern metrics, a Kruskal-Wallis test was used to determine the level of marking size differences between ‘mean bird’ models against their own and different biomes, and between all bird models (‘mean bird’ and ‘quantile bird’) to their own biome at different spatial scales of the backgrounds.

## Results

### Prediction: plumage attributes of specialist bird species will differ in a consistent manner between biomes

Broadly, analyses of photographs of 384 individual skins from 75 species of ground-nesting birds indicate differences in plumage appearance based on biome specialisation.

### Colour and luminance metrics

As predicted, the colour metrics of the dorsal plumage patches show that there were significant differences among specialist ground-nesting bird species across biomes (Hue: H = 101.8 (6), p < 0.001. Saturation: H = 176.1 (6), p < 0.001. Luminance: H = 511.3 (6), p < 0.001). In general, the dorsal plumage patches of tropical rainforest, taiga forest, and dry forest birds were characterized by low luminance and high hue and saturation values. By contrast, the dorsal plumage patches of desert, grassland, and tundra birds exhibited high luminance and lower hue and saturation values (for full details, see Fig. 2 and Table 3).

**Figure 2.**
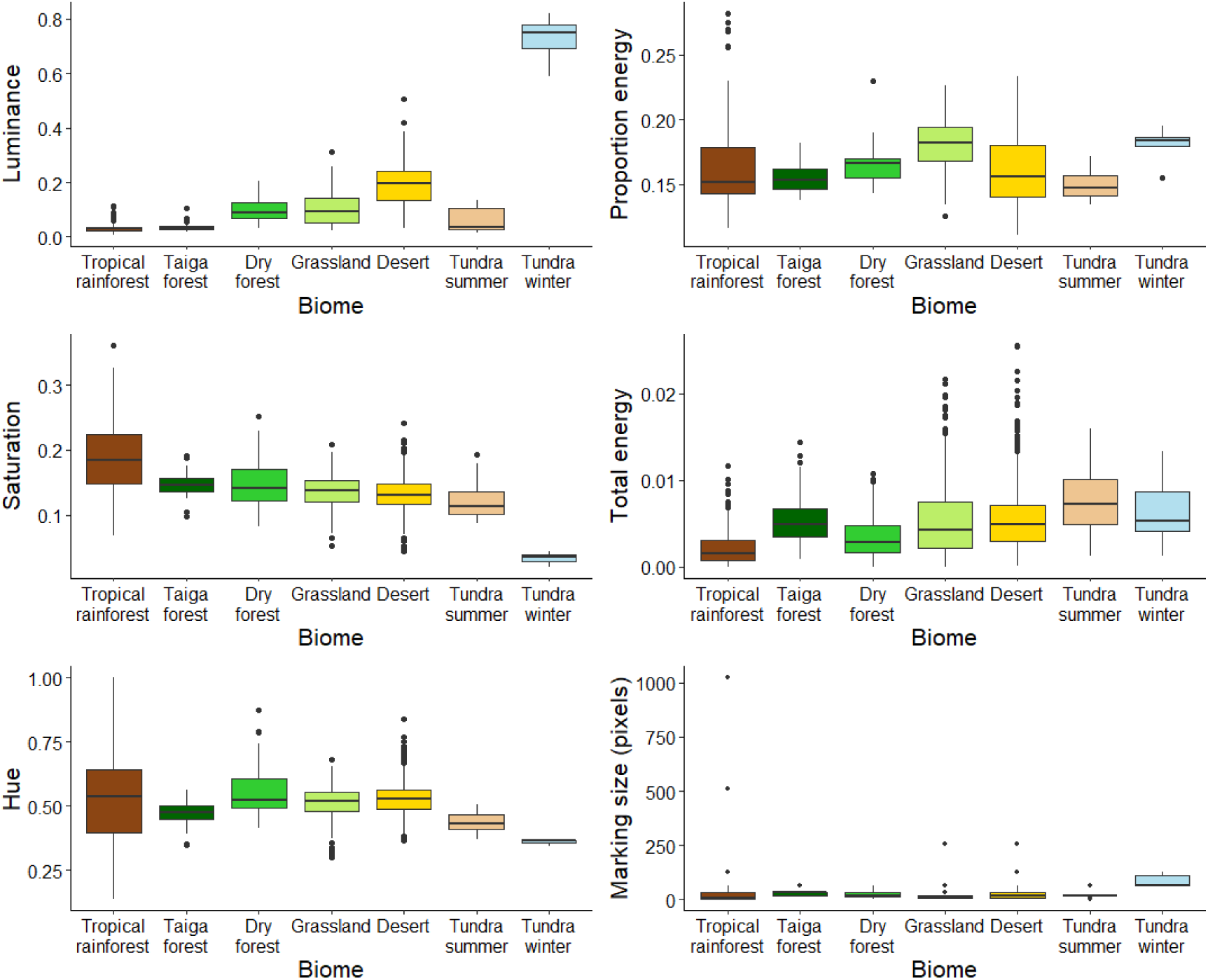
Absolute values of the dorsal plumage attributes of specialist ground-nesting bird species of different biomes. In general, the results show that specialist ground-nesting bird species differ in their dorsal plumage attributes based on biome specialisation. Specifically, tropical rainforest birds exhibited a dark, rich brown/red coloration with the lowest pattern contrast. Taiga forest birds exhibited a dark, rich brown/green coloration with high pattern contrast. Dry forest birds showed a bright, rich red/yellow coloration with moderate to low pattern contrast. Grassland and desert birds exhibited a pale, red/yellow coloration with moderate to high pattern contrast. The summer plumage of tundra birds showed a pale, brown/purple coloration with the highest pattern contrast, whereas the winter plumage displayed a pale, white appearance with high pattern contrast. Tundra winter = winter plumage. Tundra summer = summer plumage. Boxes represent interquartile ranges (IQRs) with centre lines indicating medians, whiskers represent the lowest and highest values within 1.5 × IQR, and filled circles indicate outliers.to observations within 1.5 × IQR, and points beyond the whiskers indicate outliers.

With respect to the colour metrics of the ventral plumage patches, there were significant differences among specialist ground-nesting bird species across biomes (Hue: H = 23.6 (6), p < 0.001. Saturation: H = 48.9 (6) p < 0.001. Luminance: H = 138.7 (6), p < 0.001). In general, with the exception of taiga forest, dry forest, grassland, and the summer plumage of tundra birds, the colour appearance (i.e. hue, saturation and luminance) of ventral plumage patches in tropical rainforest, desert and the winter plumage of tundra birds, was similar to that of their dorsal plumage (for full details, see Fig. 3 and Table 3).

**Figure 3.**
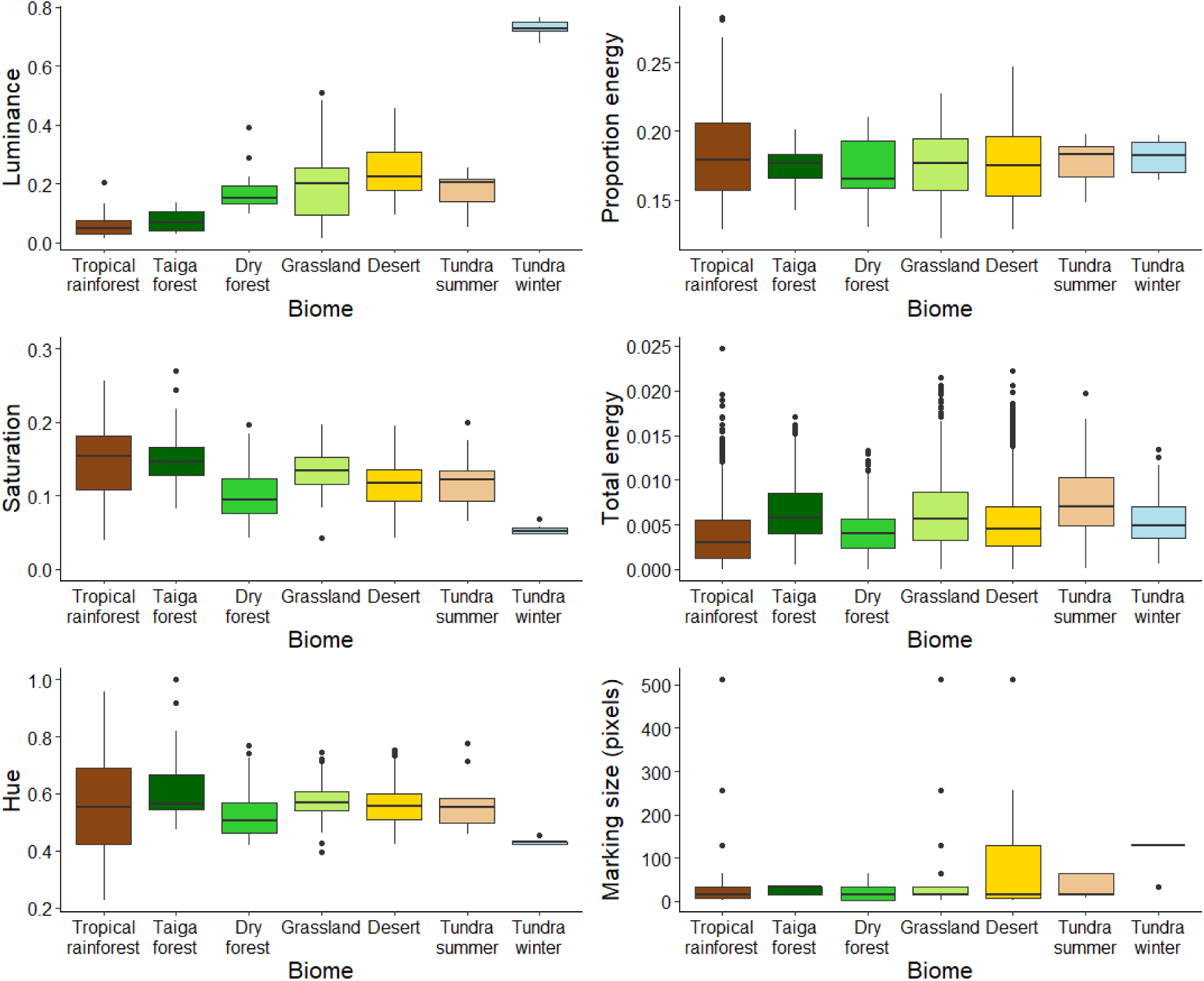
Absolute values of the ventral plumage attributes of specialist ground-nesting bird species of different biomes. In general, the results show that specialist ground-nesting bird species differ in their ventral plumage attributes based on biome specialisation. Specifically, tropical rainforest birds exhibited a dark, rich brown/red coloration with moderate to low pattern contrast. Taiga forest birds exhibited a dark, rich red coloration with high pattern contrast. Dry forest birds showed a pale, brown/green coloration with moderate to low pattern contrast. Grassland birds showed a pale, red coloration with high pattern contrast. Desert birds exhibited a pale, red/yellow coloration with moderate to high pattern contrast. The summer plumage of tundra birds showed a pale, red/yellow coloration with the highest pattern contrast, whereas the winter plumage displayed a pale, white appearance with moderate to high pattern contrast. Tundra winter = winter plumage. Tundra summer = summer plumage. Boxes represent interquartile ranges (IQRs) with centre lines indicating medians, whiskers represent the lowest and highest values within 1.5 × IQR, and filled circles indicate outliers.

### Pattern metrics

As predicted, the pattern metrics of the dorsal patches show that there were significant differences among specialist ground-nesting bird species among biomes (Total energy: H = 567.5 (6), p < 0.001. Proportion energy: T = 16.8 (6) p < 0.001. Marking size: T = 10.34 (6), p < 0.001). In general, across biomes, dorsal plumage shows mainly small dominant markings with a single maximum pattern energy peak, except in grassland birds, which display more than one distinct energy peak (for full details, see Figs. 2 and 4; Table 3).

**Figure 4.**
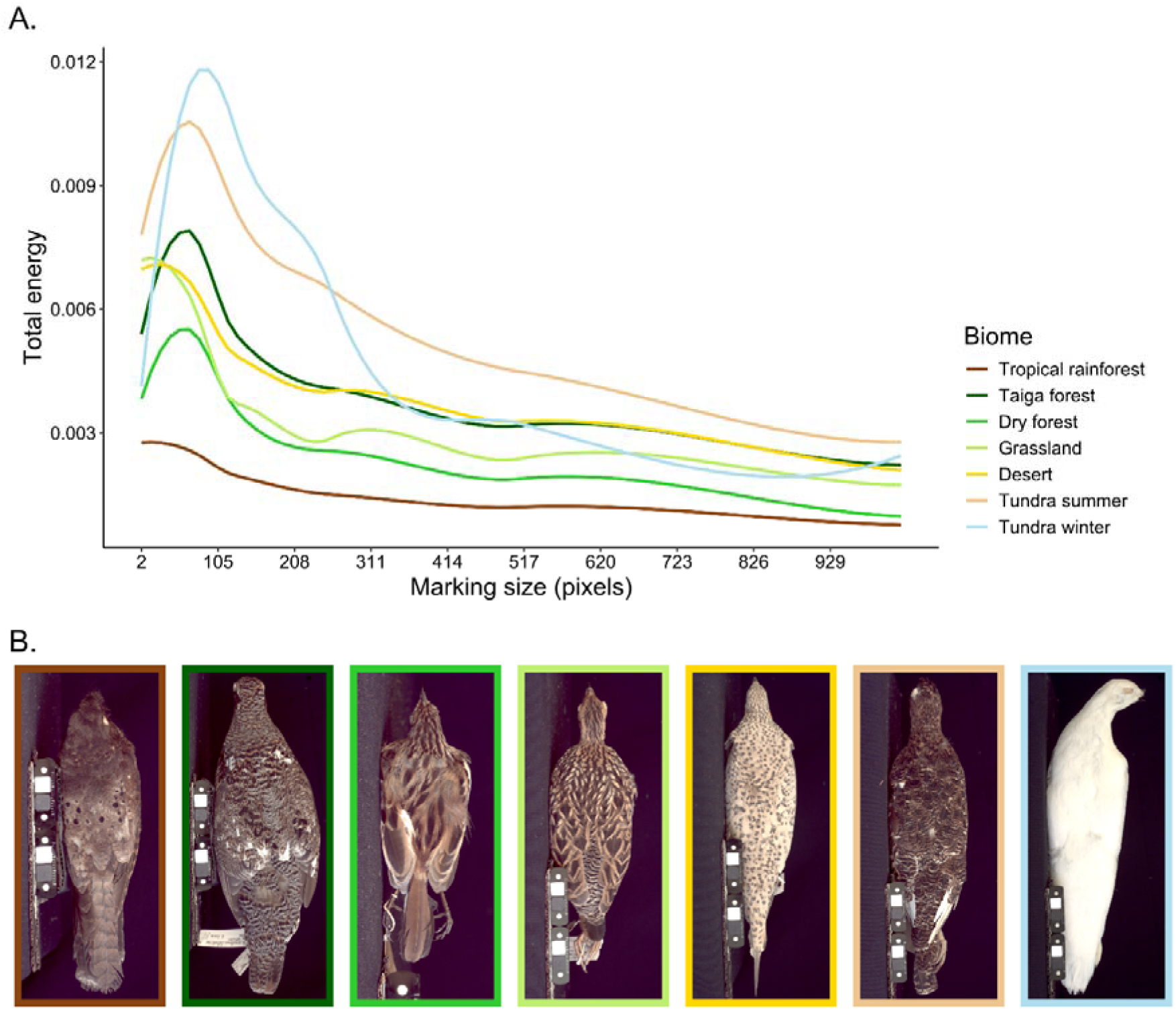
Granularity spectra from dorsal plumage patches of specialist ground-nesting bird species of biomes. A) Dorsal plumage pattern contrast of tropical rainforest, taiga forest, dry forest, desert, and tundra birds showed small marking sizes with a single, well-defined peak in pattern energy, whereas grassland birds exhibited small marking sizes reaching more than one distinctive pattern energy peak, one well-defined and a second, weaker peak occurring at larger marking sizes. B) Examples of dorsal plumage of specialist ground-nesting birds. From left to right, the species are: *Nyctiphrinus ocellatus*, *Canachites canadensis*, *Myrmorchilus strigilatus*, *Francolinus pictus*, *Pterocles coronatus*, and *Lagopus lagopus.* Tundra winter = winter plumage. Tundra summer = summer plumage.

In terms of ventral plumage, the total energy and marking size results show that there were significant differences among specialist ground-nesting bird species of different biomes for total energy (H = 207.8 (6), p < 0.001) and marking size (H = 16.4 (6), p < 0.011), but not for proportion energy (T = 0.4 (6), p = 0.861). In general, across all biomes, ventral plumage exhibited small markings that either reached a single maximum pattern energy peak, whereas birds from taiga forest, grassland, and desert birds, displayed more than one distinct energy peak (for full details, see Figs. 3 and 5; Table 3).

**Figure 5.**
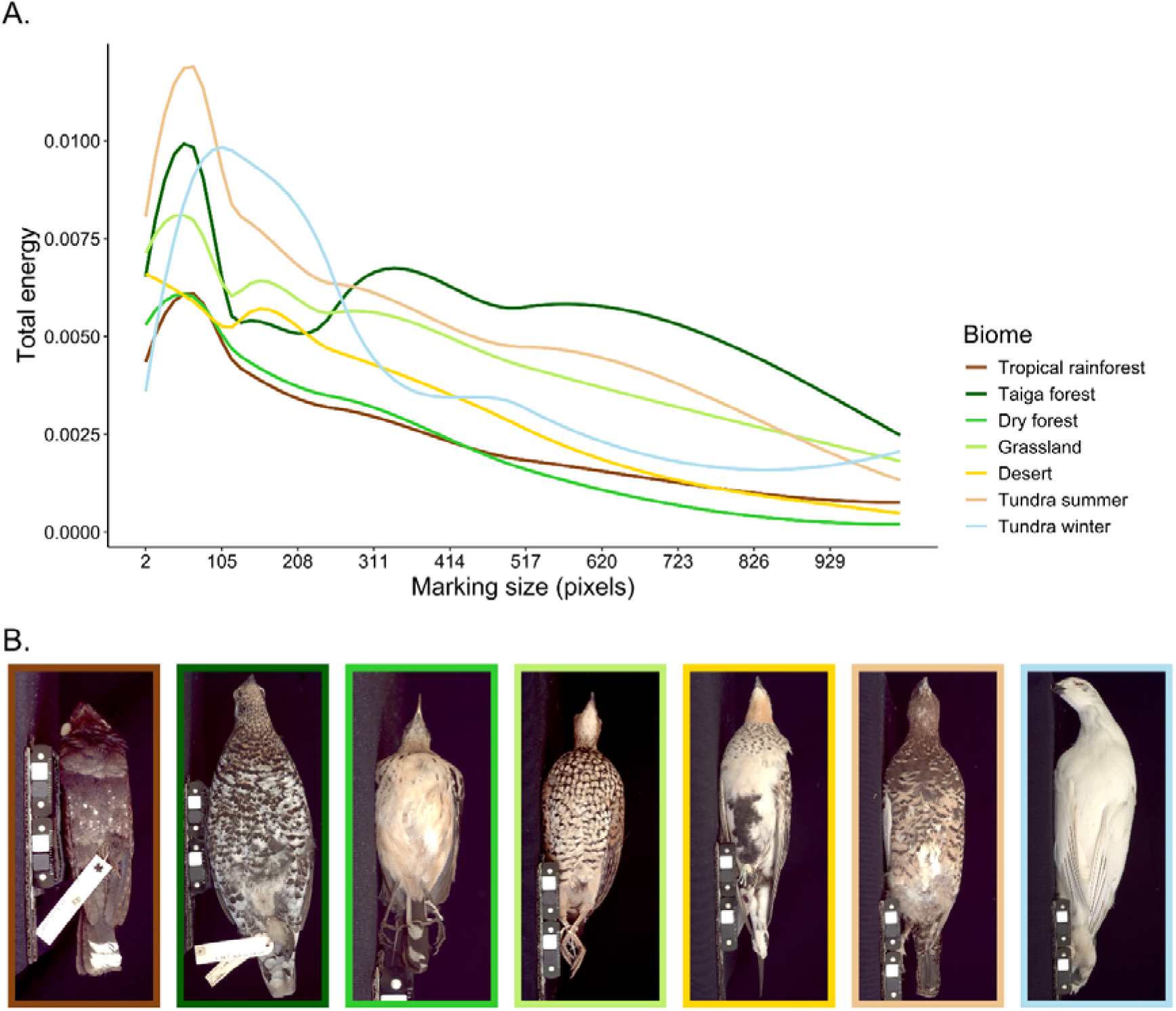
Granularity spectra from ventral plumage patches of specialist ground-nesting bird species of biomes. A) Ventral plumage pattern contrast of tropical rainforest, dry forest, and tundra birds showed small marking sizes with a single, well-defined peak in pattern energy, whereas taiga forest, grassland, and desert birds exhibited small marking sizes reaching more than one distinctive pattern energy peak, one with higher energy and a second, weaker peak occurring at larger marking sizes, particularly in taiga forest birds. B) Examples of ventral plumage of specialist ground-nesting birds (same species as in Figure 4). Tundra winter = winter plumage. Tundra summer = summer plumage.

### Prediction: bird models will achieve better camouflage against the substrate composition and the vegetation structure of their own biome than against that of a different biome

There were substantial differences between how well tropical rainforest and grassland ‘mean’ bird models matched natural backgrounds when placed in their own versus a different biome (interaction between ‘Source Birds’ and ‘Photographed Biome’: F1,86 = 49.49, p <0.001 for dorsal coloration, F1,86 = 26.11, p <0.001 for ventral coloration, F1,194 = 6.22, p = 0.013 for dorsal marking luminance, F1,194 = 1.55, p = 0.213 for ventral marking luminance, H1 = 1.83, p = 0.175 for dorsal marking size, H1 = 14.61, p <0.001 for ventral marking size). Specifically, tropical rainforest birds exhibited a better dorsal colour and marking luminance match to the temperate rainforest biome than to the grassland biome. By contrast, grassland birds showed a better dorsal and ventral colour match to the grassland biome than in the temperate rainforest biome, while their dorsal marking luminance showed a similar match across both biomes. Contrary to the main predictions, with respect to ventral marking size, tropical rainforest birds exhibited a better match to the grassland biome than to the temperate rainforest biome, while grassland birds showed a better match to the temperate rainforest biome than to the grassland biome. Results were not significant for ventral marking luminance and dorsal marking size (Fig. 6).

**Figure 6.**
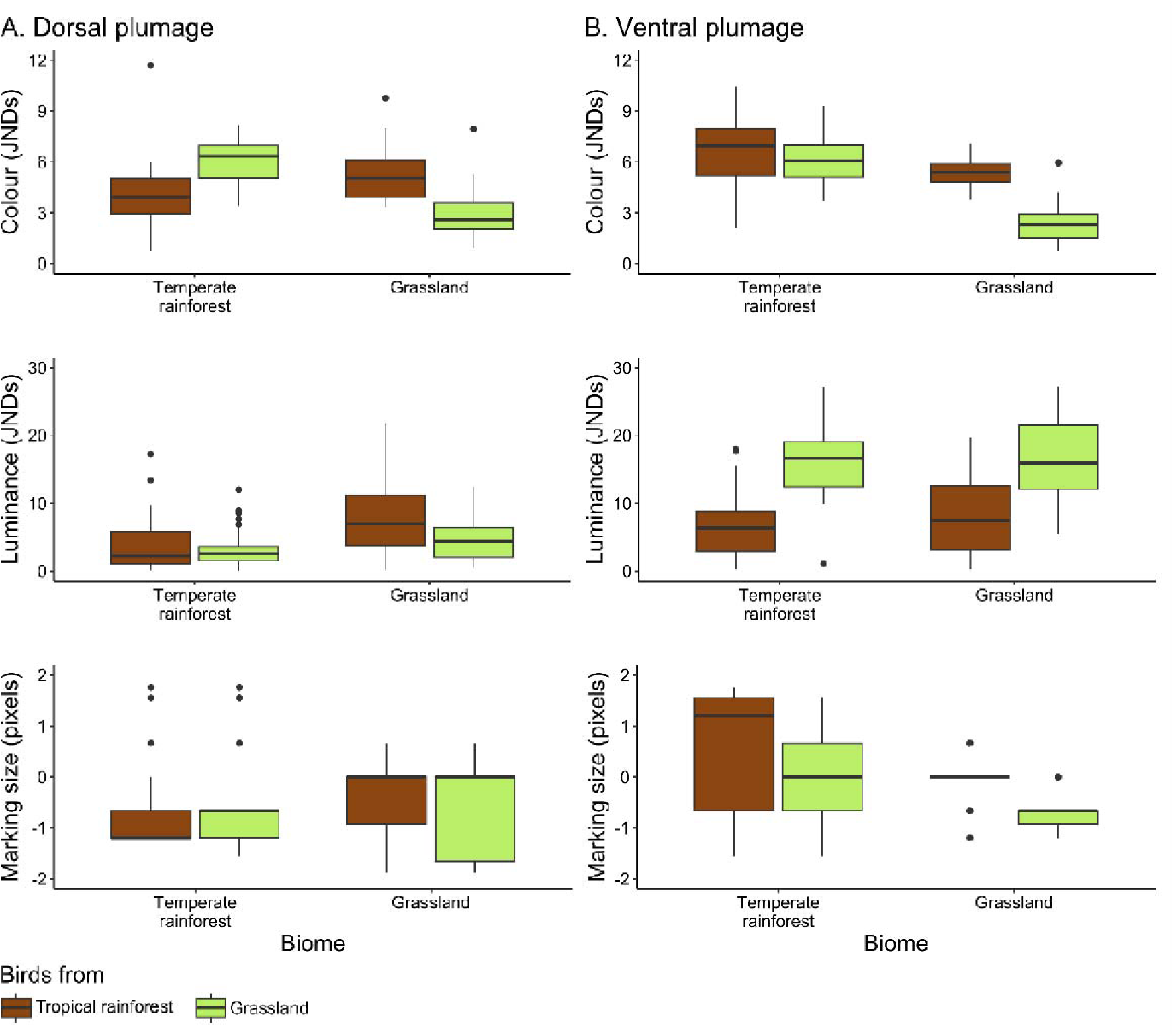
Pairwise comparisons of the matching degree of bird mean models against temperate rainforest and grassland biomes. Dorsal colour and marking luminance of tropical rainforest birds, and dorsal and ventral colour of grassland birds, show a better match to their own biome than to a different biome. Contrast in colour and marking luminance are measured in just noticeable differences (JNDs). For contrast in marking size, values of 0 represent perfect background matching, whereas values below and above 0 indicate relatively larger or smaller bird markings with respect to the background. See main text for details.

### Prediction: bird models should better match their own biome backgrounds over larger rather than smaller spatial scales

As also predicted, there were substantial differences in how well grassland and tropical rainforest bird models matched their own biome at larger spatial scales than smaller ones. Grassland bird models better matched the grassland biome at larger spatial scales compared to smaller ones (distance ID: F119.1 = 10.6, p < 0.001 for dorsal colour, F119.03 = 30.1, p < 0.001 for ventral colour, F147.17 = 3.4, p = 0.033 for dorsal marking luminance, F200.05 = 7.9, p <0.001 for ventral marking luminance, H2 = 5.5, p = 0.061 for dorsal marking size, H2 = 9.2, p = 0.009 for ventral marking size). Specifically, grassland birds showed a better match in dorsal and ventral colour at larger spatial scales compared to smaller ones. Interestingly, and contrary to the main predictions, with respect to the dorsal marking luminance, grassland birds show a better match at small spatial scales compared to large ones, while ventral marking size showed a suboptimal match across spatial scales. Results for dorsal marking size were non-significant (Fig. 7). Similarly, tropical rainforest bird models better matched the temperate rainforest biome at larger spatial scales than at smaller ones (distances ID: F120.05 = 46.3, p < 0.001 for dorsal colour, F120.02 = 23.7, p < 0.001 for ventral colour, F151 = 0.9, p = 0.396 for dorsal marking luminance, F202 = 2.8, p = 0.060 for ventral marking luminance, H2 = 14.9, p <0.001 for dorsal marking size, H2 = 30.5, p <0.001 for ventral marking size). Specifically, tropical rainforest birds showed a better match in dorsal colour and ventral marking size at larger spatial scales compared to small ones. The ventral coloration of tropical rainforest birds showed a relatively better, yet still suboptimal, match at larger spatial scales compared to smaller ones. Contrary to the main predictions, tropical rainforest birds show a better dorsal marking size match at small spatial scales compared to large ones. Results for marking luminance were non-significant (Fig. 8).

**Figure 7.**
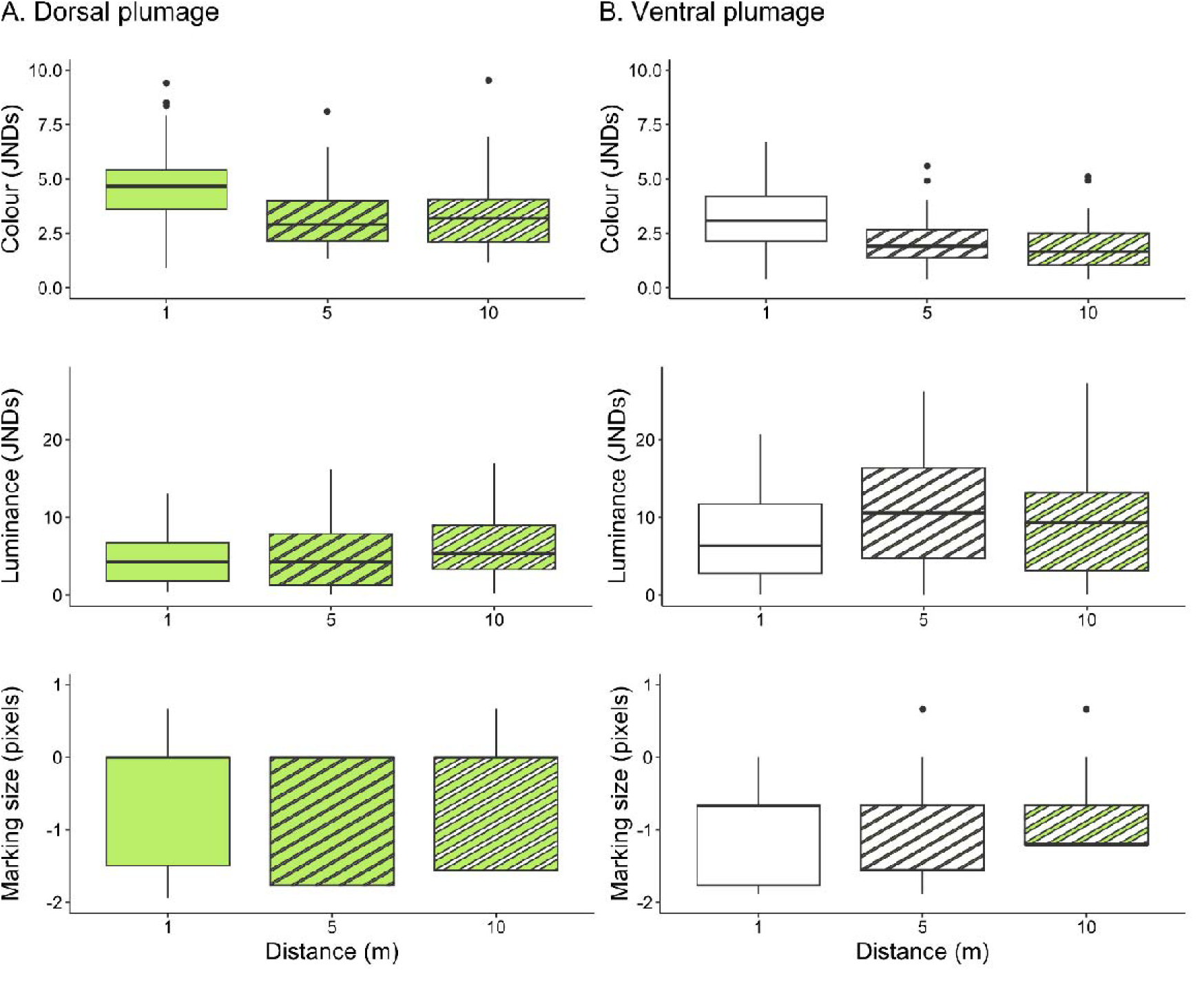
Differences of dorsal and ventral plumage match degree of quantile grassland bird models across spatial scales of grassland biome. Specialist ground-nesting birds show a better match at larger spatial scales than at smaller scales, particularly for dorsal and ventral colour. Same legend as figure 6. See main text for details.

**Figure 8.**
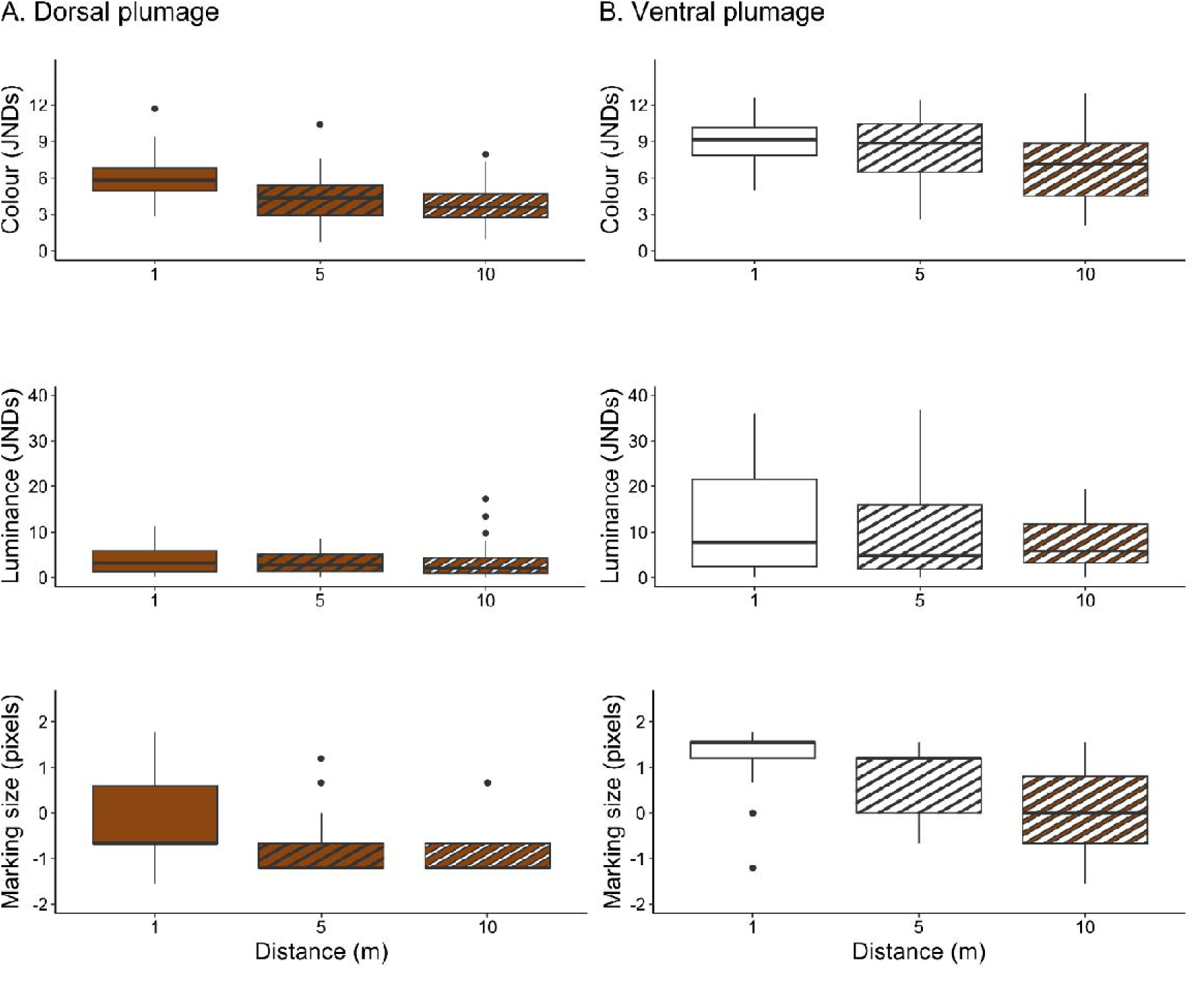
Differences of dorsal and ventral plumage match degree of quantile tropical rainforest bird models across spatial scales of temperate rainforest biome. Specialist ground-nesting birds show a better match at larger spatial scales than at smaller scales, particularly for dorsal colour and ventral marking size. Same legend as figure 6. See main text for details.

## Discussion

This study tackles the question of how animal camouflage works across different biomes through the visual system of their key predators and the potential distances at which detection may occur. Here, we tested for phenotype-environment associations in specialist ground-nesting birds found in different biomes, and how this provided camouflage. First, we found that the appearance of ground-nesting bird species differs across biomes, owing to associations between plumage colour and pattern attributes and the general characteristics of the substrate composition and vegetation structure of their biome. Second, we found that colour of grassland bird models, as well as the dorsal colour and the marking luminance of tropical rainforest bird models, are better matched to these general characteristics from their own biome than to a different one. Finally, we found that the colour of grassland bird models, and the dorsal colour and the ventral marking size of tropical rainforest bird models, show a better match against the broad characteristics of their own biome across large spatial scales, rather than at more variable and patchy smaller scales. Especially regarding dorsal colour, this likely confers a camouflage match to the wider appearance of their biome, as perceived by raptors from scanning and detection distances.

Differences in plumage appearance among birds likely arise from consistent differences in key environmental features, such as the influence of vegetation structure on the amount of light reaching the ground. Among forest biomes, distinct amounts of light reaching the ground (Endler, 1993) are associated with differences in plumage luminance (Figs. 2 and 3; Table 3). Birds with the darkest plumage occur in tropical rainforest and taiga forest, both characterised by closed canopies that strongly reduce light levels at the ground. Rainforest bird species inhabiting the understory in particular exhibit dark dorsal plumages that match the low light conditions (Endler & Théry, 1996; Gomez & Théry, 2007). Conversely, likely due to the semi-closed canopy structure, the comparatively higher plumage luminance of dry forest birds indicates an association with the higher light levels reaching the ground.

In biomes with open vegetation structure, besides being associated with high light conditions, differences in plumage luminance are closely linked to sunlight reflection from the main substrate composition (i.e. albedo). Indeed, it is possible to associate the albedo along a decreasing gradient from the highest levels observed in snow, followed by desert and grassland substrate (Table 2; e.g. Cooke, 2012) with the highest plumage luminance in tundra-winter birds, followed by high levels in desert birds and moderate to high levels in grassland birds, respectively (Figs. 2 and 3; Table 3). In a similar relationship, Rosenblum (2006) found a clear background matching between the brightness values of desert lizards and the dark and white soil backgrounds. Differences in plumage luminance may also reflect variation in background albedo associated with the general characteristics of their biome. In the tundra biome, as the background albedo increases from snow-free substrate and arctic vegetation to snow (Table 2; e.g. Belke-Brea *et al.,* 2020), changes in the plumage luminance of tundra bird species indicate an adjustment to match the sunlight reflection of their substrate composition (Figs. 2 and 3; Table 3). Adjustments in appearance to match background albedo have also been reported in sargassum crabs (*Portunus sayi*), which likely serves a camouflage function (Russell & Dierssen, 2018). In summary, the dorsal and ventral plumage luminance of ground-nesting birds appears to follow predictions based on vegetation structure in forest biomes, whereas in more open biomes it is associated with light conditions and albedo determined by both substrate composition and vegetation structure.

**Table 2.**
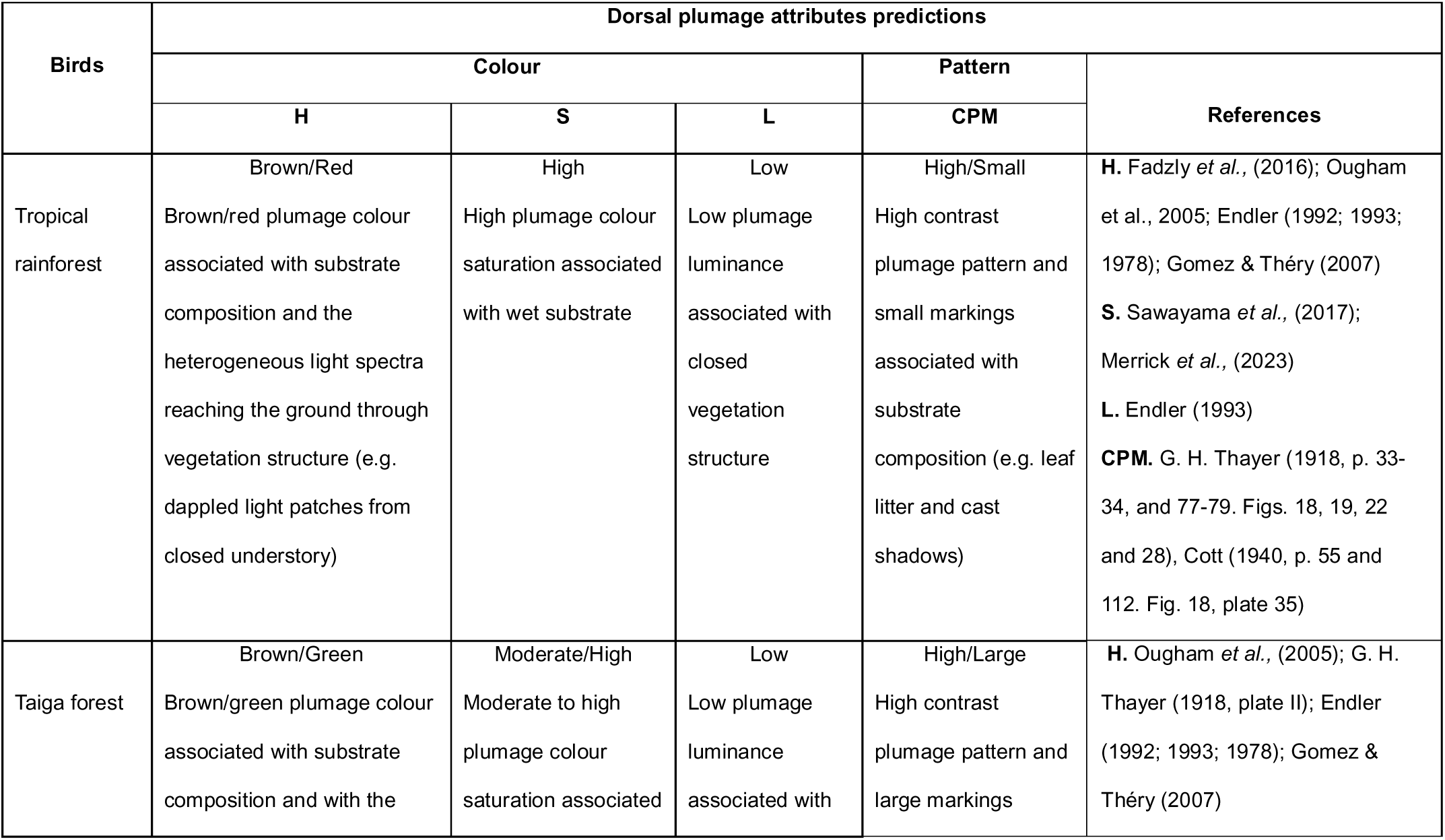
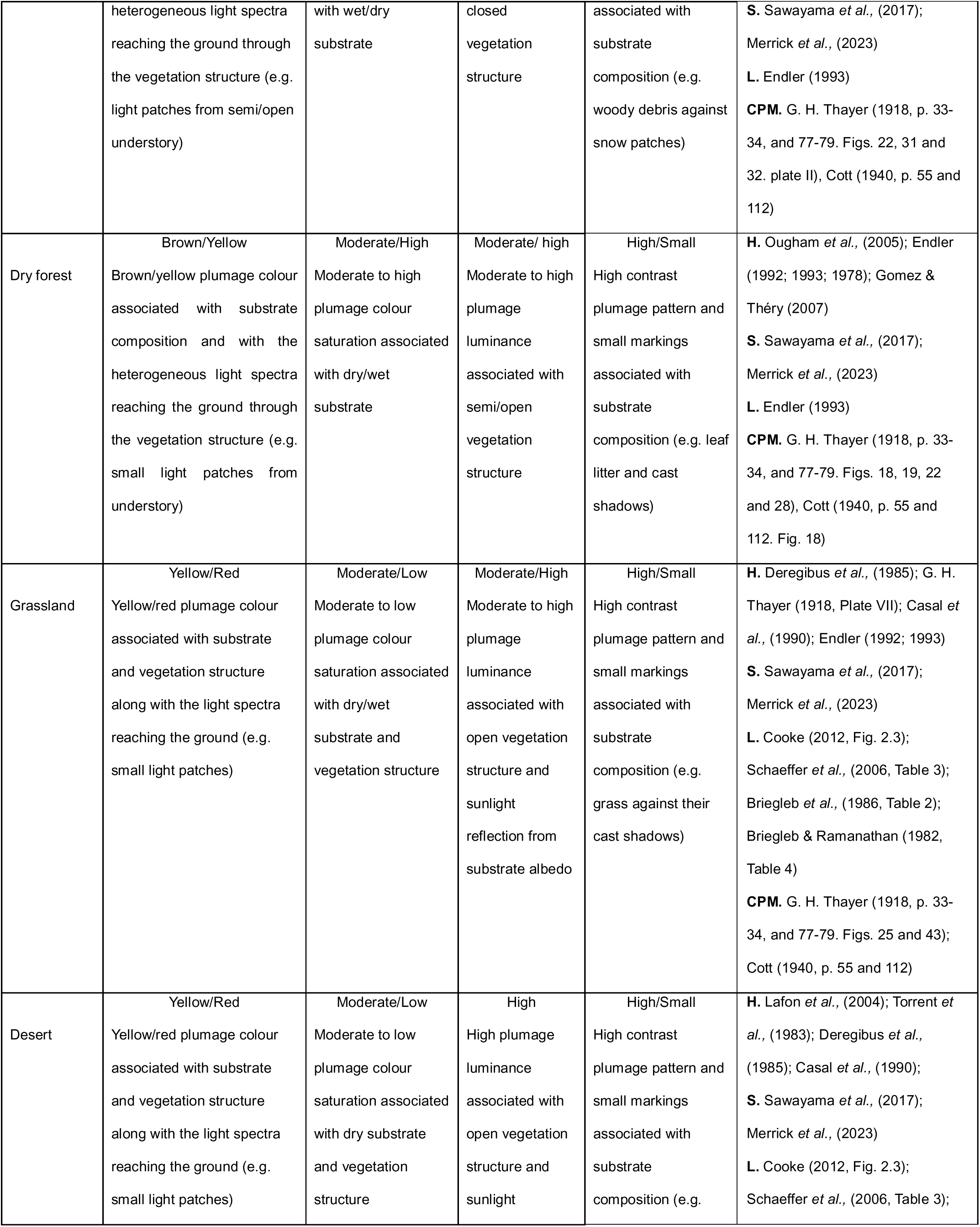
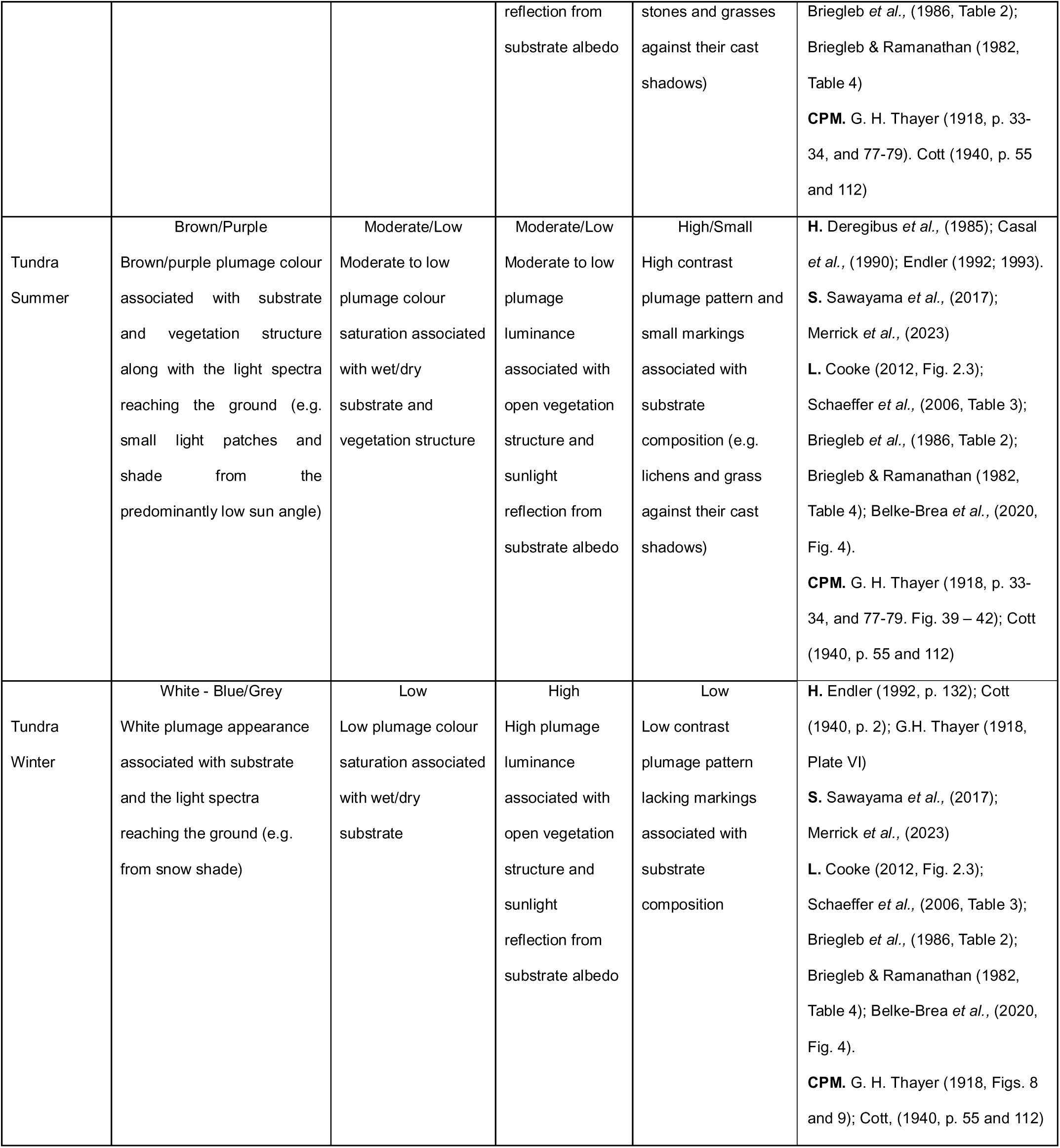
Predictions for dorsal and ventral plumage attributes of specialist ground-nesting birds of biomes, in relation to general characteristics of substrate composition and vegetation structure. The predictions in this table are based on descriptions from the literature regarding the general characteristics of substrate composition and vegetation structure previously identified for each biome of interest (see Table 1). Predictions for plumage attributes were based on the general characteristics of substrate composition and vegetation structure on (I) plumage coloration (e.g. plumage colour associated with brown/red leaf litter) along with the light spectra for plumage coloration, (II) the amount of light reaching the ground, as well as the sunlight reflectance (i.e. albedo) for plumage luminance, (III) visual perception of wetness for plumage colour saturation and (IV) environmental descriptions in camouflage artwork for plumage colour, pattern contrast and marking size (see references column). Plumage attributes are coded as follows: H = hue (colour tone), S = colour saturation, L = luminance, and CPM = contrast pattern and marking size. For tundra birds, tundra summer refers to their summer plumage and tundra winter to their winter plumage.

**Table 3.** Dorsal and ventral plumage attributes of specialist ground-nesting bird species of different biomes. Colour metrics: H = hue (colour tone), S = colour saturation, and L = luminance. Pattern metrics: TE = total energy, MS = marking size, and PE = proportion energy. Colour and pattern appearance: indicates the overall appearance of specialist ground-nesting birds based on the colour and pattern metrics (for full details, see Figs. 2 and 3. For tundra birds, tundra summer refers to their summer plumage and tundra winter to their winter plumage.

| Dorsal plumage attributes |  |  |  |  |  |  |  |
| --- | --- | --- | --- | --- | --- | --- | --- |
| Bird species | Colour metrics |  |  | Pattern metrics |  |  | Colour and pattern appearance |
|  | H | S | L | TE | MS | PE |  |
| Tropical rainforest | Highest, along with substantial variability | Highest, along with substantial variability | Lowest | Lowest | Small | Small | Among the darkest and richest plumage coloration, brown/red in tone, with low overall pattern contrast and peak energy concentrated in small, dominant markings. |
| Taiga forest | Moderate to low | High | Lowest | High | Small | Small | Among the darkest plumage colorations, rich brown/green in tone, with high pattern contrast and peak energy concentrated in small markings, tending toward small, dominant markings. |
| Dry forest | High | High | Moderate to high | Moderate to low | Small | Small | Bright, rich red/yellow plumage coloration with moderate to low pattern contrast, and peak energy concentrated in small, dominant markings. |
| Grassland | Moderate to high | Moderate | Moderate to high | Moderate to high | Small | Small | Pale, red/yellow plumage coloration with moderate to high pattern contrast, reaching two distinctive energy peaks: a well-defined peak concentrated in small, dominant markings and a second, weaker peak occurring at larger marking sizes. |
| Desert | Moderate to high | Moderate | High | High | Small | Small | Pale, red/yellow plumage coloration with high pattern contrast, and peak energy concentrated in small, dominant markings. |

| Tundra summer | Low | Moderate to low | Moderate to low | Highest | Small | Without a distinct dominance | Pale, brown/purple plumage coloration with the highest pattern contrast, and peak energy concentrated in small markings but without a distinct dominance of marking size. |
| --- | --- | --- | --- | --- | --- | --- | --- |
| Tundra Winter | Lowest | Lowest | Highest | High | Large | Large | Palest white plumage with high pattern contrast, and peak energy concentrated in small markings but dominated by larger markings. |
| **Ventral plumage attributes** |  |  |  |  |  |  |  |
| Bird species | Colour metrics |  |  | Pattern metrics |  |  | Colour and pattern appearance |
|  | H | S | L | TE | MS | PE |  |
| Tropical rainforest | Highest, along with substantial variability | Highest, along with substantial variability | Low | Moderate to low | Small | - | Among the darkest and richest plumage coloration, brown/red in tone, with moderate to low pattern contrast, and peak energy concentrated in small markings. |
| Taiga | High | High | Low | High | Small | - | Dark, rich red plumage coloration with high pattern contrast, reaching two distinctive energy peaks: a well-defined peak concentrated in small markings and a second, weaker peak occurring at larger marking sizes. |
| Dry forest | Moderate to low | Moderate to low | Moderate to high | Moderate to low | Small | - | Pale, brown/green plumage coloration with moderate to low pattern contrast pattern, and peak energy concentrated in small markings. |
| Grassland | High | High | High, along with substantial variability | Moderate to high | Small | - | Pale, red plumage coloration with high pattern contrast, reaching two distinctive energy peaks: a well-defined peak concentrated in small markings and a second, weaker peak occurring at larger marking sizes. |
| Desert | Moderate to high | Moderate | High | Moderate to high | Small, along with substantial variability | - | Pale, red/yellow plumage coloration with moderate to high pattern contrast, reaching two distinctive energy peaks: a well-defined peak concentrated in small markings and a second, weaker peak occurring at larger marking sizes. |
| Tundra summer | Moderate to high | Moderate | Moderate to high | Highest | Small | - | Pale, red/yellow plumage coloration with the highest contrast pattern, and peak energy concentrated in small markings |
| Tundra winter | Lowest | Lowest | Highest | Moderate to high | Small | - | Palest white plumage with moderate to high pattern contrast, and peak energy in small markings. |

These biome background characteristics also appear to determine plumage coloration. Indeed, in tropical rainforest, taiga forest, and dry forest birds, variation in plumage coloration is likely associated with differences in the brown/yellow pigments of leaf litter, under humid or dry conditions (i.e. higher or lower saturation, respectively), as well as light spectra reaching the ground (Figs. 2 and 3; Table 2). As such, a plausible case of ground-nesting bird association with the general characteristics of their substrate involves variation in leaf litter composition, shaped by the vegetation structure of different forest biomes. Similarly, in more open biomes, variation in plumage coloration is closely linked with these biome background characteristics in the predominant dry ground conditions. For example, in desert biomes, the yellow/red colour tones from iron oxides are widespread in the arid soils, even in suspension in the air (Table 2; Lafon *et al.,* 2004; Torrent *et al.,* 1983), to which the red/yellow plumage hue and saturation of desert birds is strongly associated (Fig. 2; Table 3). In a similar relationship, in tundra biome, the high concentration of anthocyanin pigments in arctic plants (Oberbaueri & Starr, 2002) and purplish light spectra from the shaded vegetation owing to the predominant low sun angle (see Endler 1992; 1993) might explain the brown/purple plumage coloration in tundra-summer birds. Overall, the plumage coloration and saturation of bird species of this study are strongly associated with the general characteristics of the substrate composition and vegetation structure of their biome.

Regarding pattern contrast, the high contrast pattern of tundra, grassland and desert birds are likely linked to their substrate composition, including cast shadows produced by the open vegetation structure, through mechanisms of background matching and disruptive coloration (Figs. 2 and 3; Table 3). For example, Chiao *et al.,* (2009) demonstrated that as the spatial scale and contrast of the substrate patterns increases, cuttlefish (*Sepia officinalis*) adjust their body patterns from uniform to mottle and disruptive appearances. Interestingly, as pattern contrast and spatial scale increase from winter to summer biome backgrounds, the change from a nonpatterned appearance to the highest contrast pattern in tundra bird species indicate an adjustment to match their substrate composition and shadows in combination with disruptive coloration (Figs. 4 and 5. Table 2). For the white winter plumage of tundra birds, the high contrast pattern likely results from the luminance interplay between different feather layers, as well as light reflected by the black box in which the bird was photographed. In either case, rather than being a methodological artifact, this pattern may reflect structural feather characteristics that match the shadowed pattern and light spectra reflections from the snow (Table 2). Similarly, in desert and grassland environments, disruptive appearance appears to be associated with background elements and their cast shadows (Sherbrooke 2002; Pizzigalli *et al.,* 2020) as these biomes may exhibit substrates with relatively homogeneous appearance (e.g. similar appearance between bare ground and the surrounding elements). In such context, shadows cast by background elements are strongly associated with differences in pattern contrast, marking size variability, and energy peaks in desert and grassland birds.

With respect to forest biomes, although high contrast plumage patterns were predicted, only taiga forest birds exhibited it (Figs. 2 and 4; Table 2). Despite the overall substrate composition with high contrast patterns of forest biomes, compared with the dense understory vegetation of tropical rainforests and dry forests, the semi-open understory of taiga forests may be insufficient to break up the bird body shape from the perspective of their main predators, such as the goshawk (*Accipiter gentilis*). In summary, background matching and putative disruptive coloration are linked to biomes where both the bird’s body shape and the high contrast patterns of the substrate composition are exposed by the open vegetation structure.

The better match of bird models against to their own biome reveals an avian camouflage specialisation to the general characteristics of substrate composition and vegetation structure (Fig. 6). This is particularly evident in tropical rainforest birds, where a better dorsal colour and marking luminance match against the temperate rainforest biome indicates a camouflage specialisation to the general substrate composition and ambient light caused by complex vegetation structure of rainforest biomes.

On this theme, when viewed from a greater distance and across different biome backgrounds, the better match of grassland and tropical rainforest bird models should hinder detection by matching the wider appearance of their own biome (Figs. 7 and 8). This may be because from large distances, heterogeneous backgrounds such as leaf litter turn into a more visually uniform appearance (e.g. Stamp & Wilkens, 1993); a similar effect occurs in grassland, where, at a distance, individual grass structures acquire a more general yellow/green appearance compared to short distances (e.g. Fig. 1). Ground-nesting birds might need to match the wider appearance of their biome, possibly because the specific place where they are most vulnerable to at longer-distance detection becomes more unpredictable. Similarly, when shore crabs (*Carcinus maenas*) are more likely to be found across habitat types, the dark green phenotype is more effective at preventing or delaying detection across all examined habitats (i.e. mudflats, mussel beds, and rock pools), rather than being most effective in any single one, suggesting a generalist camouflage strategy (Nokelainen *et al.,* 2019). Taken together, our results may indicate that ground-nesting birds exhibit a generalist strategy that matches the wider appearance of several habitat types within their biome, potentially compromising optimal matching at the more variable and smaller patches, decreasing the risk of detection from distances at which their predators, particularly raptors, are searching for them.

Despite the lack of phylogenetic control in our study, it is unlikely that the results would be affected by the relatedness of species, given that the study involves different genera and virtually all year-round ground-nesting bird species of specific biomes (e.g. *Lagopus* from the tundra). Future work should test the effectiveness of phenotype-environment matching to avoid detection at large spatial scales and how different camouflage types emerge and function under ecologically realistic encounter probabilities. It should also investigate the survival value of phenotype–environment matching across backgrounds that share general characteristics; for instance, tropical rainforest birds may also have a similar survival probability in biomes that share a similar vegetation structure such as temperate rainforest. Migratory ground-nesting birds or species that move between biomes are particularly good candidates to test these ideas. Phenotypes associated with large spatial scales appear to be widespread in nature, providing a valuable opportunity to investigate how camouflage types and strategies evolve and function to reduce detection against general background characteristics.

## Acknowledgements

This work is dedicated to my parents, Fernando Medel and Conchita Hidalgo, who have always supported and encouraged me in my academic journey and passion for nature.

This study was supported by the National Agency for Research and Development (ANID) of Chile, through the Becas Chile scholarship programme [scholarship number 72170372], awarded to Javier Medel to undertake studies and research at the University of Exeter, UK.

## Data availability statement

The data underlying this study will be deposited in a publicly accessible repository and made available upon publication of this article.

