## Supporting Information for "Phenotype-environment matching in ground-nesting birds across and within large-scale biomes"

**^2^** ***Department of immunology, genetics and pathology, Uppsala University, Uppsala, Sweden***

Table S1. Specialist ground-nesting bird species of biomes and sample size.

Table S2. Structure of the sampling design.

Figure S1. Example of selection of plumage colour, marking luminance and size in museum bird skins.

Figure S2. Bird models matching the colour, marking luminance and size of museum bird skins.

Figure S3. Selection of background areas across different spatial scales of biomes.

**Table S1. Specialist ground-nesting bird species of biomes and sample size.** Museum bird skin species photographed and the sample size per biome. Five museum bird skin specimens were photographed for each species. For tundra biome, eight specimens per species were photographed, five for the plumage appearance during the summer season and three for the winter appearance. Despite the small sample size for tundra and taiga forest, most of the bird species that match the selection criteria were photographed which means practically all year-round ground-nesting bird species of both biomes*.*

| Biome | Bird skins sample size | Bird species sample size | Ground-nesting bird species |
| --- | --- | --- | --- |
| Tropical rainforest | 80 | 16 | *Antrostomus sericocaudatus, Arborophila campbelli, Caprimulgus batesi, Eurostopodus archboldi., Eurostopodus papuensis, Francolinus lathami,* *Gactornis enarratus, Galloperdix bicalcarata, Guttera plumifera, Lophura edwardsi, Lophura inornata, Lophura ruffa, Nyctiphrynus ocellatus, Nyctiphrynus rosenbergi, Ptilorrhoa carulescens, Scolopax saturata.* |
| Taiga forest | 30 | 6 | *Dendrogapous fuliginosus obscurus, Falcipenis canadensis, Falcipenis falcipenis, Falcipennis franklinii, Tetrao urogallus, Tetraro urogalloides.* |
| Dry forest | 35 | 7 | *Caprimulgus clarus, Cinclosoma castanotum, Cinclosoma punctatum, Drymodes brunneopygia, Myrmorchilus strigilatus, Philortyx fasciatus, Caprimulgus donaldsoni.* |
| Grassland | 110 | 22 | *Afrotis afroides, Certhilauda semitorquata, Chordeiles pusillus, Coturnix coromandelica, Coturnix delegorguei, Cursorius temminckii, Emberiza calandra, Eupodotis caerulescens, Eupodotis senegalensis, Francolinus africanus, Francolinus gularis,* *Francolinus pictus, Lissotis hartlaubii, Lophotis ruficrista, Lophotis savilei, Pternistis swainsonii, Pterocles gutturalis, Pterocles quadricinctus, Synoicus adansonii, Sypheotides indicus, Tympanuchus cupido, Tympanuchus pallidicinctus.* |
| Desert | 105 | 21 | *Alaemon alaudipes, Ammoperdix griseogularis, Ammoper dixheyi, Callipepla gambelii, Callipepla squamata, Caprimulgus aegyptus, Caprimulgus eximius, Caprimulgus nubicus, Chlamydotis undulata, Cinclosoma cinnamomeum, Cursorius cursor exsul, Cursorius rufus, Eremopterix nigriceps, Pterocles burchelli, Pterocles coronatus, Pterocles senegallus, Pterocles lichtensteinii, Pterocles personatu, Syrrhaptes paradoxus, Syrrhaptes tibetanus, Systellura decussata.* |
| Tundra | 24 | 3 | *Lagopus lagopus., Lagopus leucorus., Lagopus mutus captus* (All terrestrial arctic tundra bird species). |

**Table S2.** **Structure of the sampling design.** Within each of the two studied biomes (Temperate rainforest and Grassland), bird models were photographed at 3 different ' locations ' and at 3 ' sites ' per location (n = 9 sites per biome). Within each site in both studied biomes, two mean bird models, each reflecting the mean colour, marking luminance and marking size of tropical rainforest or grassland birds were photographed. Within each site in the Grassland biome, also 4 quantile bird models reflecting the quantile colour, marking luminance and marking size of grassland birds were photographed. Similarly, 4 quantile bird models representing the quantile colour, marking luminance, and marking size of tropical rainforest birds were photographed at each site of temperate rainforest biome. Every bird model photographed at each site was also photographed at three distances (i.e. one metre, five metres, and ten metres) to assess the potential effects of the spatial scale at which a predator may search for the target bird. Within the temperate rainforest biome, the three different locations chosen were each in a different altitudinal habitat: one location within each of the high mountain forest, middle mountain forest and lower mountain forest. Within the Patagonian grassland biome, the three different locations chosen were each in a different lowland grassland habitat: one location within each of shrubland, yellow grass steppe (in dry grounds) and green grass steppe (in humid grounds). This selection was made with the intention to sample representative background characteristics of each biome. H.M.F = high mountain forest. M.M.F = middle mountain forest. L.M.F = lower mountain forest. Shrubland = grass and shrubs. Yellow = steppe with yellow grass appearance (in dry grounds). Green = steppe with green grass appearance (in humid grounds).

|  | **Temperate rainforest** biome | | | **Grassland** biome | | |
| --- | --- | --- | --- | --- | --- | --- |
|  | Locations | | | Locations | | |
|  | H.M.F | M.M.F | L.M.F | Shrub | Yellow | Green |
| **Tropical rainforest** ‘mean’ bird models (1 model photographed at each site, at 3 distances) | 3 sites | 3 sites | 3 sites | 3 sites | 3 sites | 3 sites |
| **Grassland** ‘mean’ bird models (1 model photographed at each site, at 3 distances) | 3 sites | 3 sites | 3 sites | 3 sites | 3 sites | 3 sites |
| **Tropical rainforest** ‘quantile bird models (4 models photographed at each site, at 3 distances) | 3 sites | 3 sites | 3 sites | - | - | - |
| **Grassland** ‘quantile’ bird models (4 models photographed at each site, at 3 distances) | - | - | - | 3 sites | 3 sites | 3 sites |

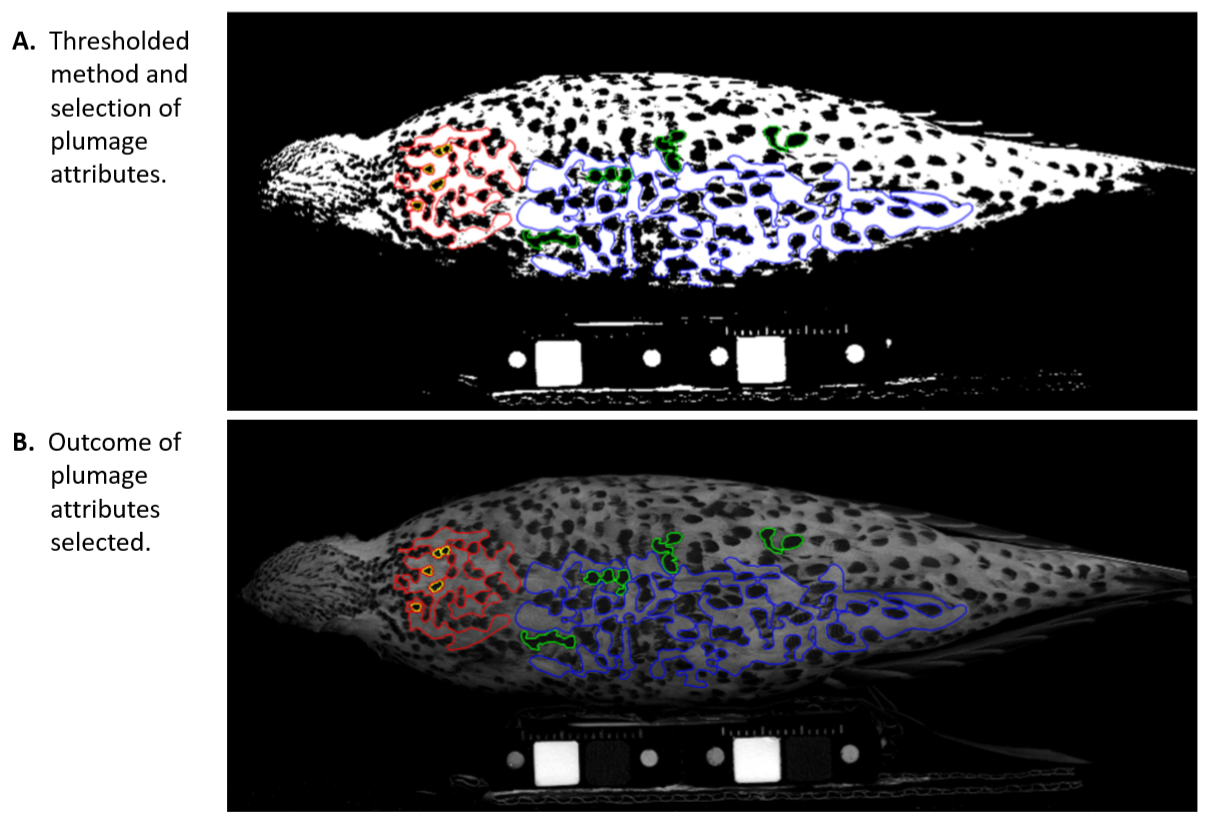

**Figure S1.** **Example of selection of plumage colour, marking luminance and size in museum bird skins.** A) Thresholded method and selection of plumage attributes: a multispectral image thresholded in a binary format, facilitating detection and selection of plumage colour, marking luminance and size attributes. B) Outcome of plumage attributes selected: the original multispectral image with the outcome of colour and marking attributes selected. The different colour polygons show plumage attributes selected for mantle and wing patches. The red polygon encloses the plumage colour, and the yellow polygons enclose marking attributes, both on the mantle patch. The blue polygon encloses plumage colour, and the green polygons enclose marking attributes, both on the wing patch. These images show an example of colour and marking selection of a female specimen of spotted sandgrouse (*Pterocles senegallus*), a specialist ground-nesting bird of desert biomes.

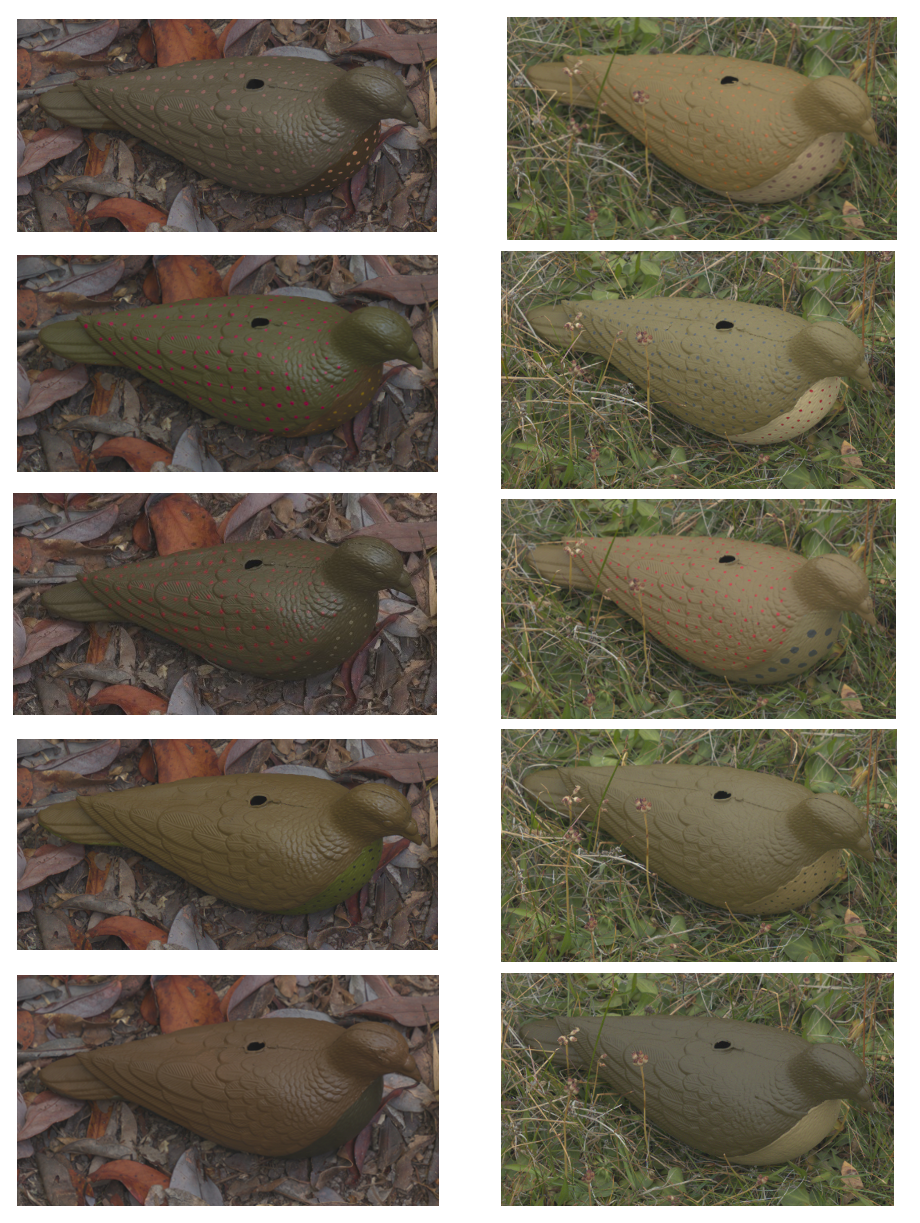

A. **Tropical rainforest bird models	B. Grassland bird models**

**Figure S2. Bird models matching the colour, marking luminance and size of museum bird skins.** A) Tropical rainforest bird models: this shows all the bird models that match the colour, marking luminance and size of specialist ground-nesting bird species of the tropical rainforest biome. B) Grassland bird models: this shows all the bird models that match the colour, marking luminance and size of specialist ground-nesting bird species of the grassland biome. For both, tropical rainforest (A) and grassland (B), bird models, from top to bottom: bird matching the 75% quantile of plumage colour, marking luminance, and marking size values from the analysis of museum specimens; followed by the 65% bird model; mean plumage match; a 35% quantile match; and a non-patterned bird treatment matching the 25% quantile.

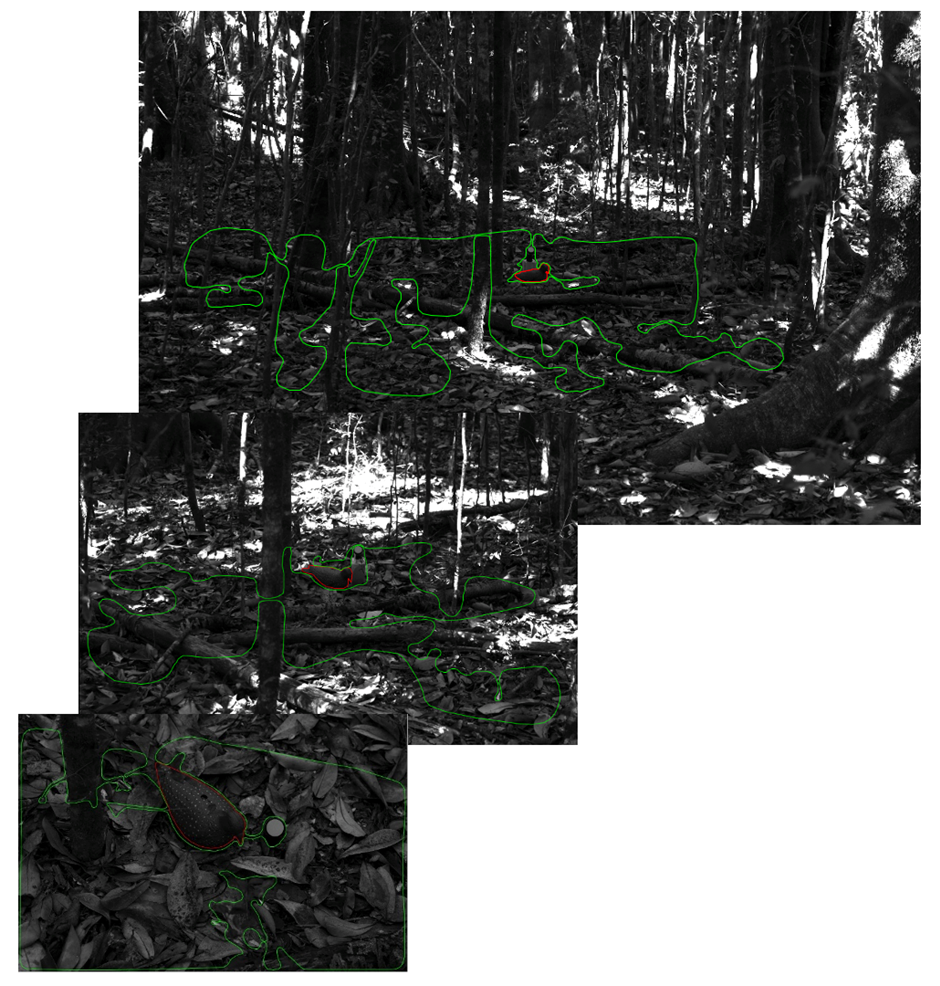

**Figure S3. Selection of background areas across different spatial scales of biomes.** The green polygon shows the background selection of different multispectral images taken at one metre twenty-five centimetres (A), five metres (B) and ten metres (C). Across different spatial scales, the background selection includes the background that surrounds the bird model and progressively encloses different background areas. The background selection excluded the dappled light conditions, the bird model (i.e. highlighted in red), areas not in focus and the grey standard. This example shows the background selection across different spatial scales of the temperate rainforest biome for a tropical rainforest quantile bird model (i.e. sixty-five percent treatment).
